# Fuel-driven catalytic molecular templating

**DOI:** 10.64898/2026.02.18.706517

**Authors:** Merry Mitra, Rakesh Mukherjee, Križan Jurinović, Thomas E. Ouldridge

**Affiliations:** Department of Bioengineering and Centre for Engineering Biology, Imperial college London, London SW7 2AZ, UK

## Abstract

Catalytic molecular templating, wherein a copolymer molecule serves as a sequence-specific template to propagate genetic information to a daughter copolymer, is fundamental to cells. Templating underlies DNA replication, RNA transcription and protein translation, underpinning the molecular basis of heredity, evolution, and biological function, and allowing staggering complexity to arise from simple building blocks. It has hitherto been challenging to emulate templating without highly evolved enzymes, largely due to product inhibition of catalytic turnover, which is a major challenge for templated dimerization and prohibitive for longer products. We present an enzyme-free DNA-based templated dimerization reaction enabled and controlled by a fuel strand that actively displaces the product from the template only once dimerization is complete, overcoming product inhibition. We systematically investigate design variants to optimise catalytic turnover, and demonstrate information propagation through the action of distinct templates that assemble specific products from the same pool of building blocks. We also show that the fuel represents an input by which the templating can be controlled, allowing the coupling of catalytic turnover to the output of upstream DNA circuitry.

**TOC Graphic:** 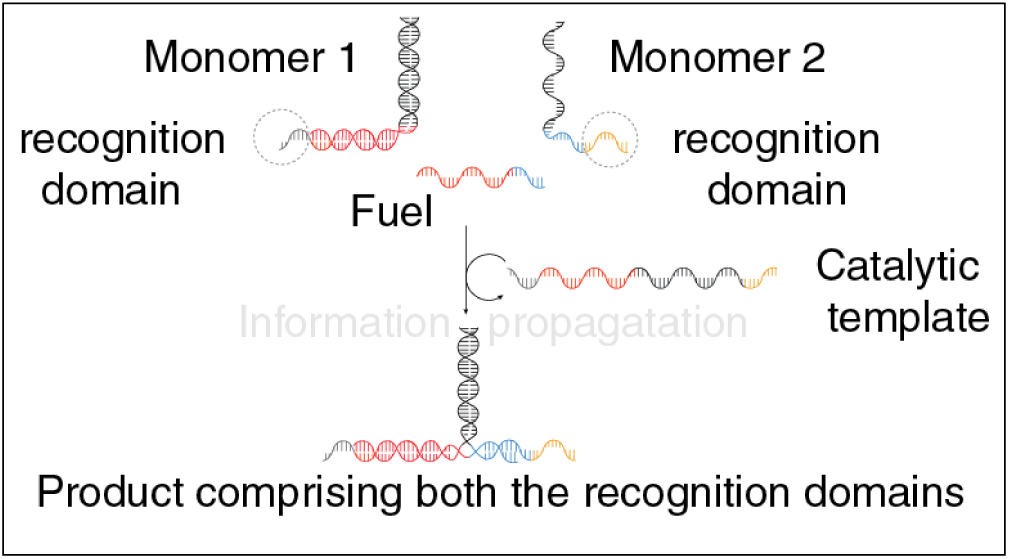

## INTRODUCTION

Molecular templating is a fundamental principle in natural biochemistry, facilitating the controlled assembly of sequence-defined macromolecules through specific recognition interactions. In biological systems, templated processes underline the core mechanisms of the central dogma, encompassing DNA replication, RNA transcription, and protein translation.^1^ In these processes, a copolymeric template strand encodes sequence-specific information that is accurately transferred to an assembling daughter strand via highly selective molecular recognition interactions.

Templating enables the biosynthesis of a vast array of functional macromolecules - most notably, tens of thousands of distinct proteins in a single organism - from a limited set of monomers, such as the 20 canonical amino acids. Without templating, this feat would be impossible, as amino acids lack the specificity required to direct the selective formation of defined protein structures from the vast space of possible polypeptide products.^2^ The information in the template circumvents this problem by enabling the formation of specific polypeptide sequences that then fold into functional proteins.

Importantly, the final product of templating must decouple from the template to allow for catalytic turnover.^3–5^ For instance, during RNA transcription, ribonucleotides base pair with a DNA template while forming a complementary RNA strand. The RNA must eventually be released, however, to become functionally active. The recognition interactions that guide sequence selection therefore do not stabilize the final product; instead, they facilitate the formation of a metastable intermediate that ultimately resolves into the detached, functional RNA.^6^ This transient binding behavior is essential to the purpose of templating.^7,8^ If templates could not be reused, a separate, sequence-specific template would have to be created for each product. Making this template from scratch would be as complex as building the product itself, defeating the main advantage of using templates.^4,5,9^

In nature, highly evolved enzymes and ribosomes allow effective catalytic templating to be the central paradigm for the production of molecular complexity. By contrast, catalytic templating is largely unexploited in synthetic systems. This absence is a major missed opportunity: in principle, synthetic, enzyme-free templating systems capable of sequence-specific polymerization would unlock a new, biologically-inspired paradigm in chemistry.^10^ These systems could exploit template-directed recognition to enhance otherwise inefficient reactions,^11,12^ synthesize sequence-controlled polymers with novel functions,^13,14^ or generate combinatorial libraries of candidate therapeutics.^15,16^ The shortage of enzyme-free templating systems also reflects our uncertainty about the mechanisms underlying prebiotic chemistry and the origins of life.^7,17–20^

The central mechanistic challenge facing the construction of templating systems is not the accuracy of recognition, but a phenomenon known as product inhibition.^21^ Product inhibition occurs when a product binds to its catalyst with sufficient affinity to impede further catalytic cycles. If the lifetime of the product–catalyst complex is comparable to that of the substrate–catalyst complex, the product competes with incoming substrates for binding to the catalyst, compromising turnover. In cases where product dissociation is extremely rare, the catalyst becomes sequestered in an inactive state, eliminating turnover altogether. This issue is especially pronounced in the synthesis of oligomers and polymers, where cooperative binding of the growing chain leads to increased affinity for the template catalyst,^4^ but product inhibition tends to be prohibitive even for templated dimerization unless mitigated.

Some researchers have used external stimuli, such as thermal cycling, chemical gradients, or mechanical agitation, to modulate the binding affinity between product and template without preventing the initial assembly on the template.^22–25^ Although effective, these strategies introduce non-autonomous or non-chemical elements that reduce system robustness and limit integration into continuous, self-sustaining chemical reaction networks. Such dependencies stand in contrast to extant biological systems, where templated reactions proceed autonomously under chemically driven, isothermal conditions.

An alternative mitigation strategy is to divert some of the free energy of the dimerization or polymerization reaction itself into destabilizing interactions with the template, driving the product to separate from the template as it is made.^5,26–29^ Despite the promise of this approach, there remain very few distinct mechanisms for chemically-driven, autonomous templating with low product inhibition, and our control over those mechanisms is limited. For example, coupling templating to the output of upstream chemical systems – as is common in biology – is yet to be demonstrated. Moreover, the design space of such systems – wherein the assembly of products must be thermodynamically favourable but tightly controlled by the presence of catalysts – is strongly constrained, making it hard to eliminate undesired behaviour such as partial reversion of reaction steps.^5^

Here, we introduce a mechanism for enzyme-free, DNA-based templated dimerization in which product bond formation does not directly lead to unbinding from the template (as in previous approaches for mitigating product inhibition).^5,26–29^ Instead, the formation of the product on the template triggers a subsequent reaction with a DNA fuel strand that detaches the product from the template (Figure 1). This design feature provides additional control over the templating reaction, allowing it to be coupled to upstream chemical reactions via the fuel molecule. Moreover, the additional step loosens some of the constraints around the design of a system under tight catalytic control, removing the need for “clamp” motifs that detabilise the product.^5^ The resultant system is capable of at least ∼ 35 − 40 cycles of catalytic turnover per template with no evidence of template-facilitated disassembly of the final product.

**Figure 1.**
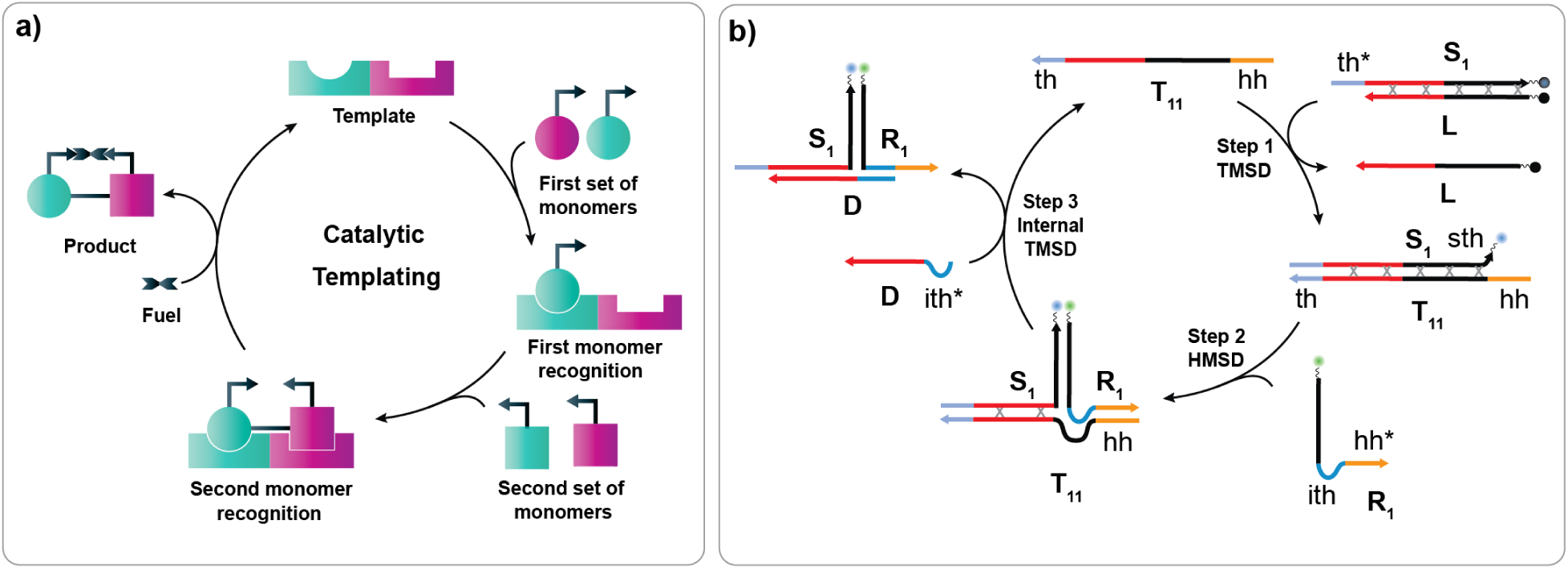
Fuel-driven molecular templating with catalytic turnover. (a) Schematic diagram of fuel-driven catalytic templating. A template binds to two monomers, each drawn from a set of alternatives according to molecular recognition interactions. Template binding accelerates formation of an initial bond between the two monomers. After this initial bonding, a third (generic) fuel molecule is incorporated into the product, stabilizing the product and separating it from the template, allowing turnover. (b) Illustration of a DNA-based realization of template-catalyzed dimerization under the control of a fuel molecule. Monomer S_1_ recognizes the template T_11_ through a short recognition domain *th* and binds to it through TMSD, releasing lock strand L (step 1). Monomer R_1_ then recognises T_11_ through a second recognition domain *hh*, binding to T_11_ and then S_1_ via HMSD (step 2). Release of the product from the template is triggered by a generic fuel strand D via internal TMSD using the internal toehold on monomer R_1_ (step 3). The inclusion of variants of both monomers, which are identical apart from their recognition domains *th* and *hh*, would allow different templates to propagate information by selective templating. The asterisk denotes a complementary domain, e.g., *th* binds with *th\**, and grey x symbols represent mismatches that provide thermodynamic driving.^30^

We first systematically characterise the individual steps of the fuel-driven catalytic templating process, optimizing its speed and exploring its catalytic turnover. We then consider a pool of competing monomers, and demonstrate that catalysts can propagate information by specifically assembling sequence-complementary products. Finally, we demonstrate how templating can be dynamically regulated through the integration of upstream DNA-based logic gates that control the availability of the fuel molecule.

## RESULTS AND DISCUSSION

### Overall design logic

Our design combines several variants of DNA strand displacement (DSD). ^5,31–35^ In DSD reactions, an invading strand displaces an incumbent from a duplex with a target strand by competing with it for base pairs. In the most common variant, toehold-mediated strand displacement (TMSD),^31^ a short single-stranded overhang on the target strand at one end of a double-stranded DNA (dsDNA) molecule – referred to as a *toehold, th* – plays a key role in controlling the kinetics of the reaction. This domain serves as a binding site for the invader, facilitating a branch migration process that ultimately leads to displacement of the incumbent strand and formation of a new duplex between invader and target. An alternative topology, in which the single-stranded *handhold, hh,* recognition domain is located on the incumbent rather than the target, is referred to as handhold-mediated strand displacement (HMSD).^35^ HMSD can be assisted by a short *secondary toehold, sth,* (1-2 nucleotide) on the target. Toeholds are not restricted to terminal positions; when located within the duplex region, they are termed *internal toeholds, ith*. Associative toeholds^34^ describe cases where the toehold binding site and the branch migration domain are contained in two different DNA strands. If these sequences are brought into close proximity during an experiment, TMSD can occur.

We combine these mechanisms in the catalytic templating cycle shown in Figure 1b. In the first step of the reaction (shown in detail in Figure 2a), the blocked monomer S_1_-L reacts with the template T_11_ via TMSD using toehold *th* to form S_1_-T_11_, revealing a single nucleotide secondary toehold *sth* on S_1_. Binding with the handhold *hh* on T_11_ and initiating the branch migration through this secondary toehold, a second monomer R_1_ reacts with S_1_ via HMSD, forming the S_1_-T_11_-R_1_ complex (Figure 2b). In this step, mismatches were eliminated to provide a thermodynamic drive to the reaction.^30^

**Figure 2.**
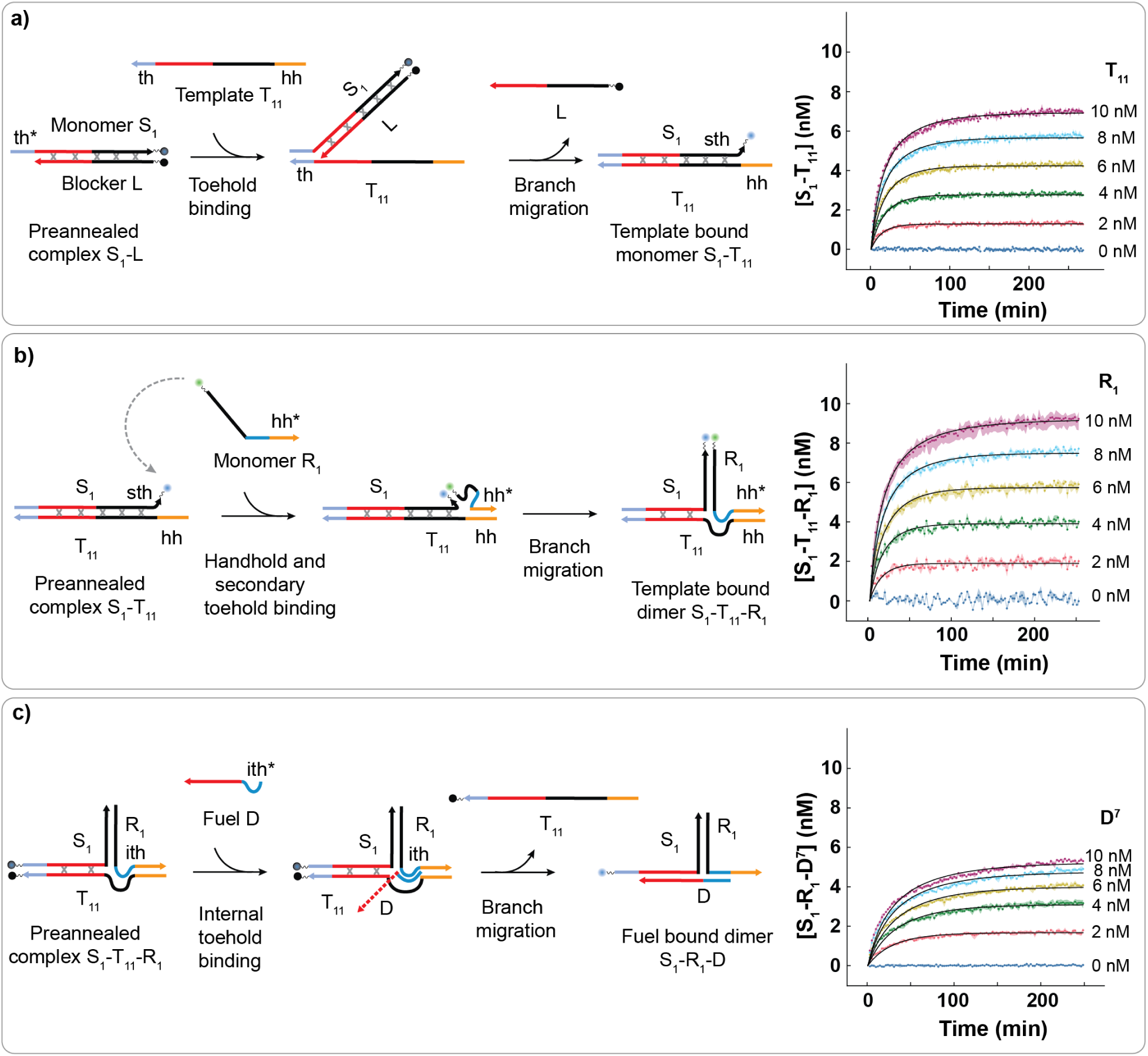
Individual experimental steps of the template-catalyzed dimerization cycle under the control of a fuel molecule. (a) Schematic of binding of S_1_ to T_11_ via TMSD in Step 1 of the catalytic cycle, and the concentration of S_1_-T_11_ against time when reacting 10 nM of S_1_-L with varying amounts of T_11_. (b) Schematic of the binding of R_1_ to S_1_ via HMSD in Step 2 of the catalytic cycle, and the concentration of S_1_-T_11_-R_1_ against time when reacting 10 nM of S_1_-T_11_ with varying amounts of R_1_. (c) Fuel-driven release of the template T_11_ to form S_1_-R_1_-D^7^ in Step 3 of the catalytic cycle, and concentration of S_1_-R_1_-D^7^ against time when reacting 10 nM of S_1_-T_11_-R_1_ with varying concentrations of D^7^. Details of how concentrations are inferred from raw fluorescence traces are provided in sections SI.3 and SI.4 of the Supporting Information, and the data are obtained from three replicas with the mean and range plotted in each case. All quoted reactant concentrations are those intended in the experimental design; pipetting errors and uncertainties in the supplied stock concentration lead to variation in the true levels, which are estimated as discussed in sections SI.3.3 - SI.3.5.

In previous work,^5^ HMSD itself was used to fully destabilise the binding of the product to the template, triggering disassociation and completing the catalytic cycle. Here, we pursue a different strategy. HMSD is used to form an initial bond between the two monomers, but the S_1_-R_1_ complex remains stably bound to the template, because HMSD halts before the interaction of S_1_ and T_11_ is sufficiently weakened. However, the formation of the dimer connects an internal toehold *ith* in the second monomer (coloured blue in Figure 1b) to a branch migration domain in the first monomer (coloured red in in Figure 1b). In this way, an associative toehold is formed. This connection allows a fuel strand D to perform a strand displacement reaction, repairing further mismatches and detaching the product from the template, completing the cycles and making the template available for catalysis in the next reaction cycle. This step is shown in detail in Figure 2c.

The net result of the intended cycle is that the two monomers S_1_ and R_1_, containing the toehold *th* and handhold *hh* recognition domains, are recognised by the template and joined together with the assistance of the fuel. The final three-stranded complex is thermodynamically favoured relative to the initial state of separated monomers and fuel, largely due to the elimination of mismatches initially present in the S_1_-L complex, but its formation is kinetically frustrated by the presence of the blocker L. The catalytic template is intended to provide a pathway for the reactions to occur.

In this process, the fuel strand functions as a generic ancillary component, engaging in interactions independent of either *th* or *hh*, or the catalyst template. Therefore, although the final product has three strands, we still classify the process as templated dimerization, as exemplified in Figure 1a, since two strands containing recognition domains are recognised by the template and joined together, and the fuel used is independent of which sequences are recognised. Although the fuel itself does not act catalytically, its role is reminiscent of the function of a helicase enzyme,^36,37^ in separating the template from the product.

As we will show, including the fuel-based step provides an orthogonal mechanism by which the reaction can be controlled. First, catalytic turnover can be modulated by the presence of the fuel strand, allowing upstream circuitry to influence templating. Second, the need for the fuel to stabilise the product means that its formation via uncatalysed “leak” reactions requires two reactions, rather than just a single blunt-ended TMSD between S_1_ and R_1_. This fact buys extra freedom when designing a system that is thermodynamically driven towards product assembly, but kinetically frustrated in the absence of a catalyst. Here, we use that freedom to avoid using “clamp” base pairs at the end of the duplex of the blocked monomer S_1_-L that are absent in the S_1_-R_1_ product. These clamps suppress leak at the expense of destabilizing the product, which contributed to the partial reversion of reaction steps seen in Ref.^5^

### Characterization of the individual reaction steps

There are many possible detailed design choices consistent with the overall design logic, including the choice of all toehold and handhold lengths, the length of the branch migration domains and the number and location of mismatches. In practice, previous experience with combining TMSD and HMSD^5,35^ suggests the use of primary toeholds *th* and handholds *hh* in the region of 6-10 nucleotides (nt), and a secondary *sth* toehold of length 1 nt. Initial testing of different toehold and handhold lengths (see Supporting Figures S2, S3 and S8), showed that a primary toehold of length of 7 nt and a handhold of length of 8 nt are suitable for our purposes. Accordingly, all further experiments were performed using this combination of |*th*| = 7, |*hh*| = 8 and |*sth*| = 1.

We elected to incorporate 5 mismatched base pairs into the initial blocked duplex to ensure that the overall reaction is sufficiently thermodynamically favourable. To minimise unintended leak reactions,^30^ these mismatches were placed at least 8 base pairs (bp) away from the end of the S_1_-L duplex, and at least 6 bp away from each other. In all experiments, we used a lock strand of length 47 nt, of which 25 nt could be replaced by R_1_ (including the elimination of 3 mismatches) and 22 nt could be replaced by the fuel D, eliminating 2 further mismatches. Since the internal displacement via an associative toehold is the least well understood process involved in the catalytic cycle, we explicitly consider variants of the fuel *D^j^* that could form internal toeholds *ith* of lengths *j* = 5, 6, 7 and 10 nt, modulated by reducing the length of D alone. Full sequences are provided in Supporting Table S2.

Figure 2, in which individual reaction steps were monitored through strand displacement-mediated quenching and unquenching of fluorophores (as illustrated in the schematic), shows that each of the individual steps in the catalytic cycle performs as required. In this case, a fuel D^7^ with |*ith*| = 7 is used for step 3 (Supporting Table S14); equivalent results for other internal toehold lengths are shown in Supporting Figure S4. Fitting simple second order reaction models to each strand displacement step, as outlined in Supporting Information SI.4 yields estimated rate constants of 1.199 × 10^5^ ± 1.039 × 10^4^*M^−^*^1^*s^−^*^1^, 1.037 × 105 ± 7.802 × 10^3^*M^−^*^1^*s^−^*^1^ and 7.363×10^4^ ± 4.727×10^3^*M^−^*^1^*s^−^*^1^ for the initial TMSD, the subsequent HMSD, and the final TMSD via an internal, associative toehold of 7 nt, respectively.

It is worth considering whether any of the reaction steps have an appreciable rate of reverse reaction - we assume they do not to perform the fits above. For steps 1 and 3, it is straightforward to test for said reversibility by mixing the two reaction products and looking for a change in fluorescence; we plot such data in Supporting Figures S5 - S7. For step one with |*th*| = 7, and for variants of step 3 with |*ith*| = 5 − 10, no significant reverse reaction was observed. For step 2, there is only one product molecule and so this procedure cannot be performed; the very low yield for |*hh*| = 7 in Supporting Figure S3 may indicate a dominant reverse reaction in that case, but the yield is much higher for the handhold length we use, |*hh*| = 8.

### Turnover in the full catalytic system

Having confirmed that the individual reaction steps proceed as expected, we now verify that mixing the monomers, template and fuel leads to templated catalytic dimerization as desired, using the protocol outlined in Supporting Table S18. In Figure 3a-d, we report the increase in fluorescence of the initially quenched fluorophore on S_1_, as 10 nM each of S_1_-L, R_1_ and D^j^ are converted into S_1_-R_1_-D by the action of the varying concentrations of template, for fuels D^j^ that form internal toeholds of length *j* = 5, 6, 7 and 10. Variants with different handhold and toehold lengths are reported in Supporting Figure S8.

**Figure 3.**
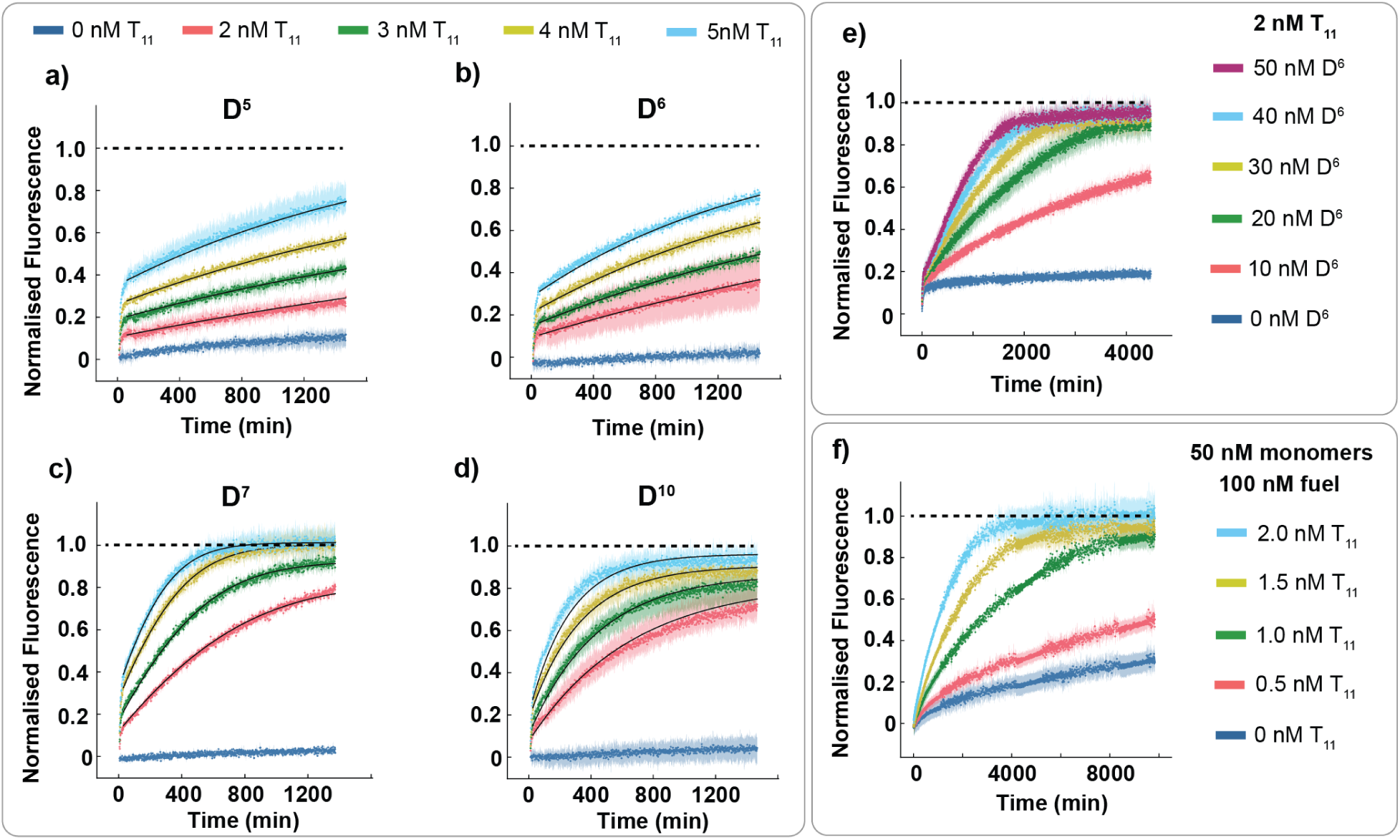
The templating mechanism exhibits effective catalytic turnover. (a)-(d) Fluorescence relative to positive control of 10 nM S_1_-R_1_-D, measured in experiments in which varying concentrations of template T_11_ were added to solutions of 10 nM each of S_1_-L, R_1_ and D^j^, in which *j* = 5, 6, 7, 10 indicates |*ith*|. Curves are fit with Equation 1 as outlined in section 4.2.1 of the Supporting Information. (e) Catalytic turnover with excess concentration of fuel D^6^. Fluorescence relative to a 10 nM S_1_-R_1_-D^6^ positive control, when 2 nM of T_11_ was added to a solution of 10 nM S_1_-L, 10 nM R_1_ alongside excess concentrations (10 - 50 nM) of D^6^. (f) Catalytic turnover with a very high concentration of monomers compared to the template. Fluorescence relative to a 50 nM S_1_-R_1_-D^7^ positive control after T_11_ at a range of low concentrations was injected into a solution of 50 nM S_1_-L, 50 nM R_1_ and 100 nM D^7^. (a)-(e) are data obtained from three replicas with the mean and range plotted in each case, whereas f is from data repeated only once; a third experiment showed a technical malfunction and is reported separately along with data at longer times in Supporting Figure S10.

We observe that for internal toeholds of length 7 and 10, the fluorescence approaches that of the positive control for all non-zero concentrations of T_11_. This behaviour indicates effective catalytic turnover. As expected for a catalyst, higher concentrations of template lead to faster reactions but a similar eventual yield. Although the number of fluorescent species in the full reaction make accurate inference of concentrations hard, an approximate quantification of catalytic kinetics can be obtained by assuming that the fractional yield of product is given by the relative fluorescence of the system compared to the positive and negative controls, as outlined in section SI.3. We then model the dynamics of this system very approximately with a Michaelis-Menten-like model

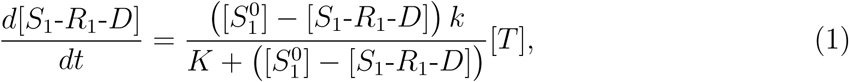

where *k* and *K* are constants. This model assumes the catalysts reach a pseudo-steady-state, and leverages the fact that both monomers and the fuel should be present at approximately the same concentration due to the particular experimental setup. Moreover, we expect each reaction step to be linear in this concentration in the low concentration regime, explaining why the unreacted concentration appears linearly in numerator and denominator.

Fits of *k*, *K* and [*S*_1_^0^] are performed as outlined in section SI.4, and illustrated in Figure 3 a-d. System behaviour is generally well-described by this model; intriguingly, initial fits for D5, D6 and D10 were deep in the unsaturated regime, meaning that only the compound variable *k/K* rather than *k* and *K* individually was meaningful. The fits plotted in Figure 3 a,b,d are therefore performed with a simpler model 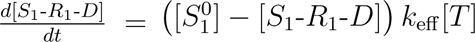, yielding *k*_eff_ of 0.00011 min^-1^nM^-1^, 0.00013 min^-1^nM^-1^, and 0.00076 min^-1^nM^-1^ for D^5^, D^6^, and D^10^, respectively. The D^7^ traces could not be fitted as well with a totally unsaturated model, with an optimal description of the data observed for *k* =0.0094 min^-1^ and and *K* = 7.77 nM, suggesting moderate saturation effects. The lack of evidence of strongly saturated kinetics suggests that, under these conditions, intramolecular rearrangement and branch migration reactions are generally fast compared to the timescale of binding.

Under these conditions, and with the modelling assumptions outlined, the D^7^ and D^10^ systems both have a maximum turnover rate of around 1 per 150 minutes for each catalyst. For D^5^ and D^6^, the fluorescence signal grows at a much slower rate, indicating less efficient fuel-driven product detachment. Indeed, for these systems at the concentrations of the experiment, the invasion of the fuel appears appears to be rate-limiting. This hypothesis is confirmed by the fact that the turnover rate can be increased by using a large excess of D^6^, as illustrated in Figure 3e.

For D^7^, by contrast, there is limited change in turnover rate when fuel concentration was increased beyond 10 nM (Supporting Figure S9), and so we conclude that the action of D^7^ is fast enough to not be rate limiting, consistent with the similar rate constants fitted to Figure 2. Equally importantly, unintended leak reactions in the absence of the template were extremely slow, showing that the assembly of the product is under strong catalytic control.

Since the catalytic turnover seen for D^7^ and D^10^ were similar, we opt for D=D^7^ as the standard internal toehold henceforth. This choice should minimise the potential for unintended internal toehold binding prior to the formation of the S_1_-T_11_-R_1_ complex, although we note that gel electrophoresis of mixtures of D and R_1_ show no evidence of premature binding even for D^10^ at high concentrations (see Supporting Figure S14). Using this design, we demonstrate that the fuel-based mechanism for templating assembly is resilient to product inhibition and capable of a high number of catalytic cycles by introducing small concentrations of template (0.5 - 2 nM) to large concentrations of monomer (50 nM) and fuel (100 nM), with the results plotted in Figure 3f. The system converges on its steady state without any dramatic slowdown due to the accumulation of product, including when product concentrations become comparable to the concentrations of first the template and then the monomer.

The high yields obtained with template concentrations as low as 1 nM are consistent with ∼ 35−40 catalytic cycles per template before the monomers were consumed. In these systems with relatively high concentrations of monomers and fuel, non-negligible leak reactions were observed. Nonetheless, catalytic acceleration of the reaction (relative to spontaneous leak) remains clear at concentrations as low as 0.5 nM of T_11_, 100-fold lower than the monomers. Note that depletion of monomers will suppress the contribution of this leak pathway in experiments with non-zero template concentration.

### Information propagation via orthogonal dimerization

Natural templates don’t merely catalyse the assembly of monomers into products, they catalyse the production of specific products from a specific sequence of monomers, thereby propagating information from parent to daughter. Here, we demonstrate that our fuel-based templating mechanism can also pass on information by selecting for specific alternative S and R monomers.

We introduce variant monomers S_2_ and R_2_ with distinct recognition domains *th* and *hh* that are orthogonal to those in the original system. Additional thymine bases were also added to the 5*^′^* end of R_2_ to make it slightly larger than R_1_, and distinct fluorophore labels were appended to the 3*^′^* ends of S_1_ and S_2_. All other domains, including those involved in branch migration and the binding to the fuel, are unchanged relative to S_1_ and R_1_ (Supporting Table S4). As a consequence, all four products S_1_-R_1_-D, S_1_-R_2_-D, S_2_-R_1_-D and S_2_-R_2_-D can form with approximately equal stability. As reported in Supporting Figure S11, the dimerization of S_2_ and R_2_ can be templated by their complementary template T_22_ in an analogous manner to S_1_ and R_1_ by T_11_. We will now demonstrate information propagation by showing that a template with the appropriate handhold and toehold domains T_ij_ can *selectively* catalyse the production of S*_i_*-R*_j_*-D.

We initially tested orthogonality by adding 2 nM of a single template T_ij_ to 10 nM each of S*_k_*-L, R*_l_*, D, for all 16 combinations of *i, j, k, l*. We plot the fluorescence relative to a positive control as the *S* monomer is liberated from the quencher-carrying blocking strand *L* for *k* = *l* = 1 and all four possibilities of *i, j* in Figure 4a. For T_11_, we see the familiar catalytic turnover and an increase towards the positive control. For T_12_, we observe an initial rapid increase of the signal, followed by a plateau at around 20% and then a gentle increase over the rest of the experiment. In this case, the 2 nM of added template recognises S_1_ and quickly reacts with it, but the non-complementary of the handholds in T_12_ and R_1_ means that the subsequent increase, relying on the reaction to occur in the absence of a handhold, was extremely slow. For T_21_ and T_22_, the increase in fluorescence was negligible. The other permutations produce an equivalent sequence-specific response, as shown in Supporting Figure S12.

**Figure 4.**
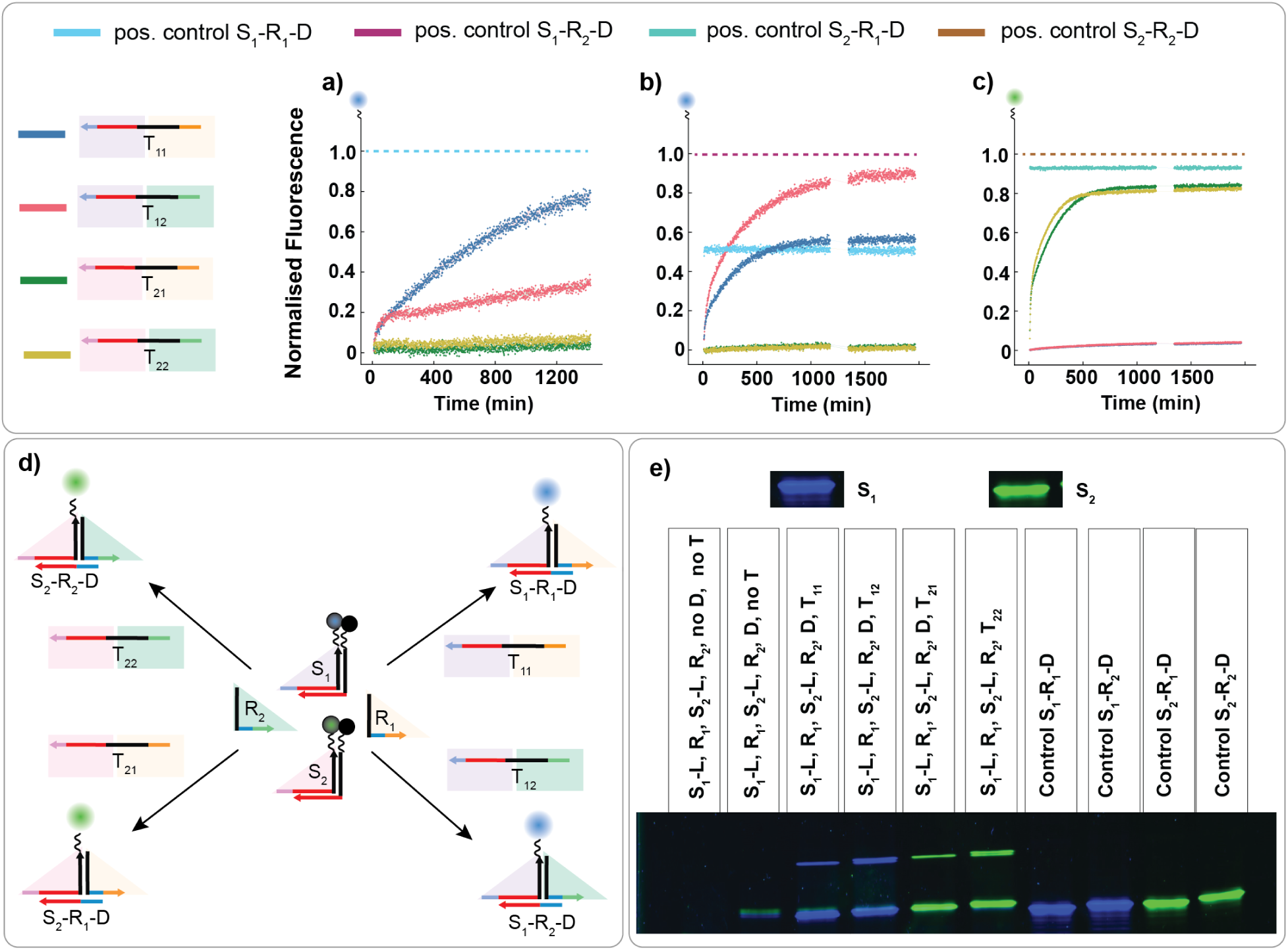
Specificity and information propagation in fuel-driven catalytic templating. (a) Fluorescence relative to a positive control of 10 nM S_1_-R_1_-D when 2 nM of template (either T_11_, T_12_, T_21_ or T_22_) was added to 10 nM each of S_1_-L, R_1_ and D. Flourescence increases as S_1_ was released from its duplex with quencher-labelled L. (b) Fluorescence in the blue channel observed after adding 5 nM of each template to a mixture of all four monomers and D, each at 10 nM, relative to positive control of 10 nM S_1_-R_2_-D (also shown is a second control of 10 nM S_1_-R_2_-D). (c) Fluorescence in the green channel observed in the same experiments as (b), relative to positive control of 10 nM S_2_-R_2_-D (also shown is 10 nM S_2_-R_1_-D). (d) Schematic illustrating the information propagation experiment from (b) and (c), in which individual templates were added to monomer pools containing of all monomers and fuel to selectively produce a matching product as in plots. (e) Gel electrophoresis image supporting the specificity of the reactions. The untrimmed gel data is reported in Supporting Figure S13.

The above results are consistent with selective catalysis. The initial TMSD step involving the S monomer is extremely specific. The specificity of the subsequent HMSD can be seen by comparing the gradients of fluorescence after the initial jump due to template binding is complete, for systems with matching and non-matching handholds. The HMSD recognition step therefore has reasonable specificity, although less than the TMSD, as evidenced by the slow but noticeable upwards drift for T_12_ in Figure 4a, and the counterparts in Supporting Figure S12. The existence of the secondary toehold and the mismatches that were eliminated during HMSD both likely contribute to the lower specificity for this step.

In Figure 4b-e, we demonstrate that this specificity can be harnessed to propagate information from template to product. As illustrated in Figure 4d, we expose each of the templates individually to the full pool of monomers. The fluorescence traces, plotted in Figure 4c-d show that S_1_ is catalytically unblocked only when T_11_ or T_12_ were added, and that S_2_ is catalytically unblocked only when T_21_ or T_22_ were added. These fluorescence signals don’t immediately tell us whether R_1_ or R_2_ has been incorporated into the product, but gel electrophoresis performed at the end of the experiments (Figure 4e), in which products containing R_1_ and R_2_ are just discernible (with the larger product migrating marginally slower) suggest that the dominant product is the correct one and that the information in the template sequence is propagated to the assembled product (the complete gel is provided in Supporting Figure S13).

### Use of the fuel to control catalytic turnover

The mechanistic requirement for a fuel molecule provides additional control over the catalytic templating process. In Figure 3e, we showed that varying the concentration of D^6^ provides an analog control over templating rate. We now show that the need for fuel allows upstream molecular logic circuits to control whether templating happens or not by dictating whether the fuel is present. The fuel is a particularly convenient handle with which to exert control over templating activity, since it can simultaneously influence many templating processes. Moreover, the fuel comprises a short single-stranded sequence of bases with only one toehold domain. Sequences like this can straightforwardly released from DNA strand displacement of fuel in a catalytic cycle using a NOT gate; input 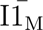 sequesters the the trigger I1_M_. In this figure, grey x symbols represent mismatched base pairs. circuits, with the other sequences used in that circuit almost independent of the fuel sequence and the templating system more broadly. In this study, we employed fuel Dn, an extended variant of fuel D^7^ that is initially complexed ancillary strand(s).

In Figure 5, we schematically illustrate designs for the generation of a fuel species through a single input DNA strand displacement gate (a YES gate); a two-input AND gate; and via a YES gate with catalytic amplification.^38–40^ We also propose the termination of fuel release by implementation of a NOT gate.^41^ These logic circuits are intended to function as regulatory checkpoints, ensuring that the fuel is only introduced under specific predefined conditions, enhancing programmability.

**Figure 5.**
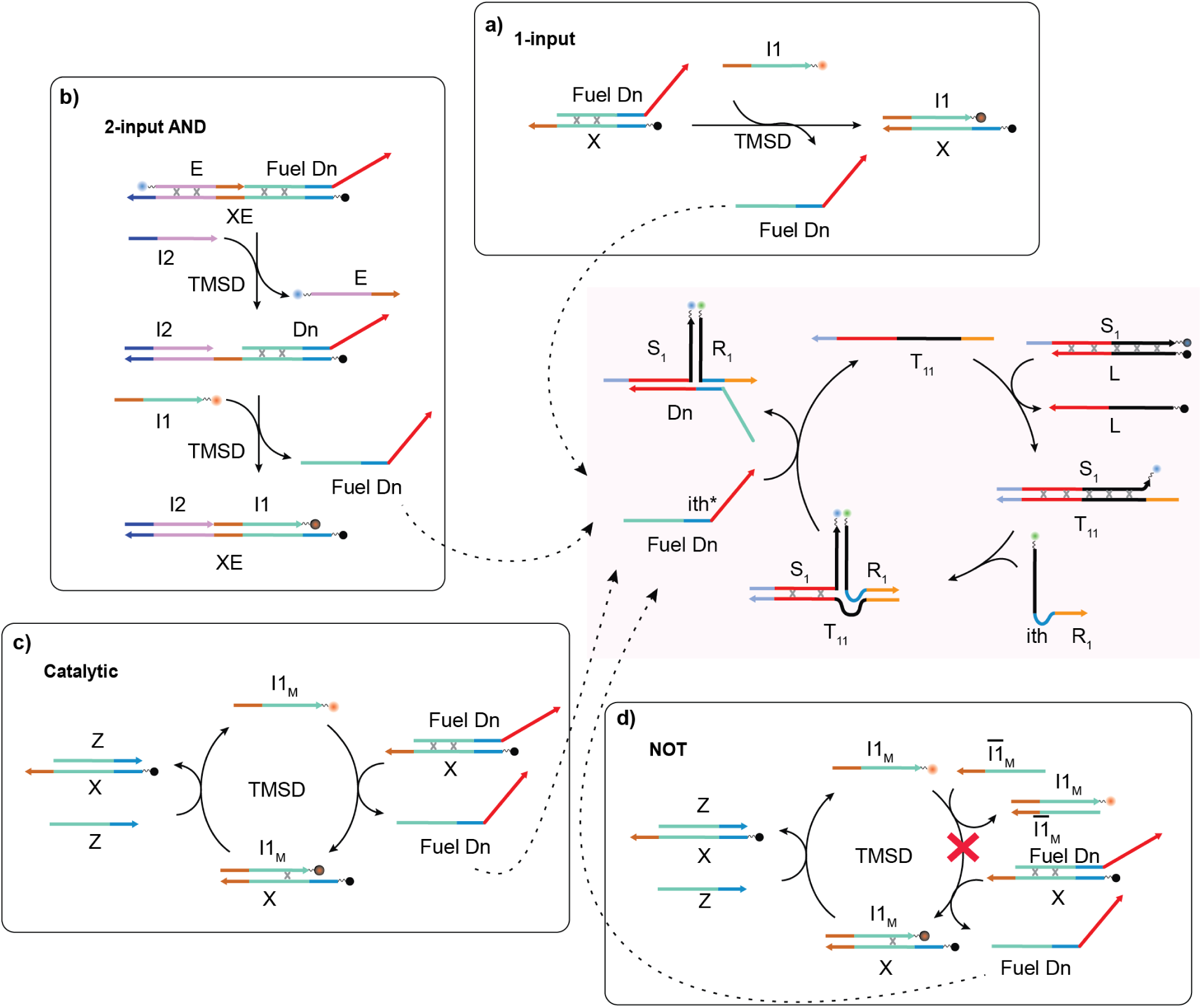
Schematic of the coupling of the templating process to upstream DNA-based logic via the fuel. The main templating cycle is the default S_1_-R_1_-D^7^ templating process considered throughout this work, with the fuel supplied as an output of the upstream circuitry. (a) 1-input YES gate in which fuel is released in response to the presence of an input I1. (b) 2-input AND gate in which fuel is released only in the presence of inputs I1 and I2. (c) 1-input YES gate in which the input signal I1_M_ is amplified via a catalytic cycle, with each I1_M_ triggering the release of multiple fuel strands D. (d) Prevention of the release

Figure 6a-d confirms that the designs in Figure 5 function as intended. In all cases, input and varying concentrations of template were injected into a solution containing 10 nM of the monomers and components of the logic circuits, and compared to controls with no input and/or no template. In this case, we used labelled R_1_ as well as labelled S_1_ and measured the resultant FRET relative to 10 nM of S_1_-R_1_-D product.

**Figure 6.**
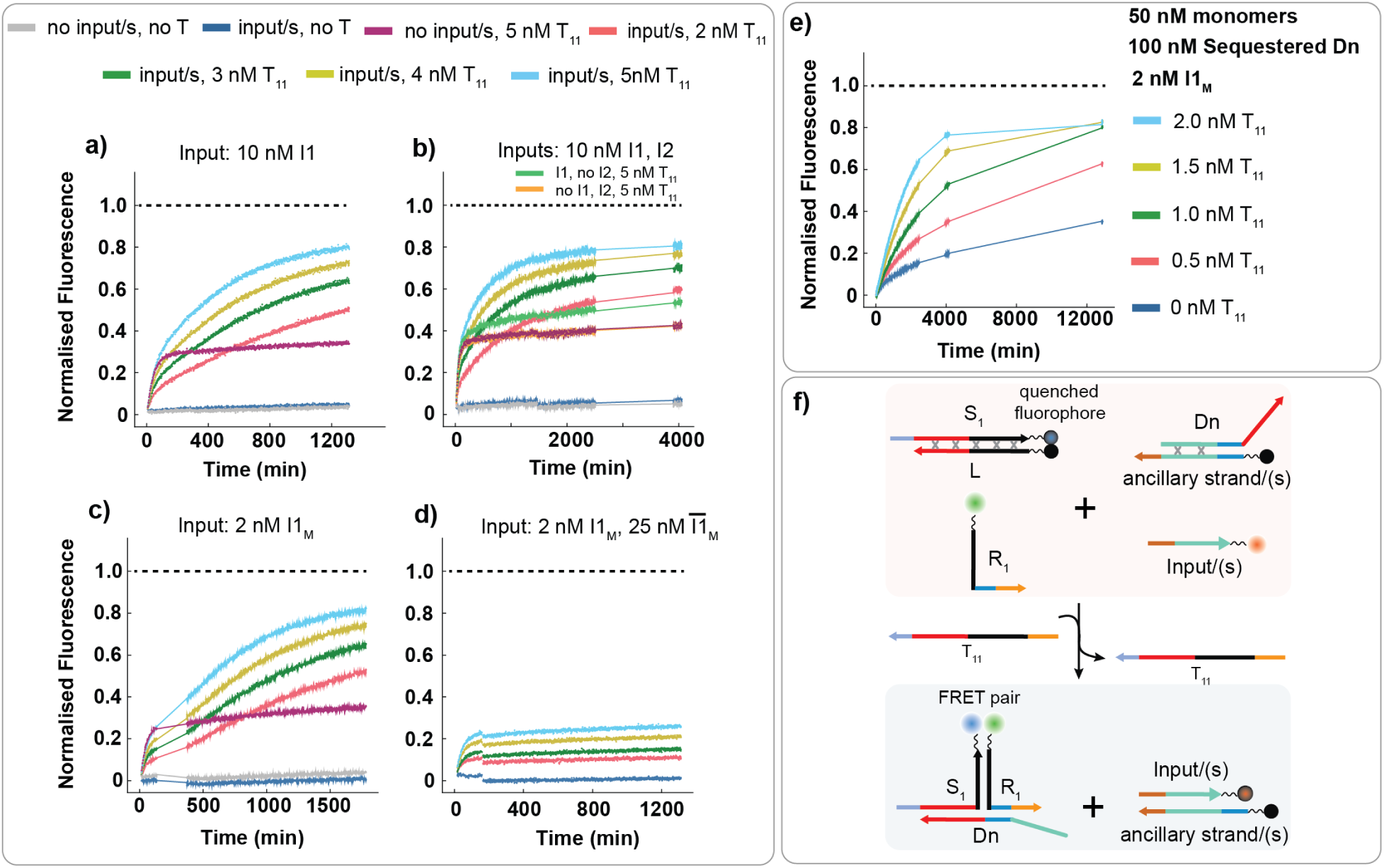
Exerting control over templating reactions through upstream logic. In each case, we report the FRET between fluorophore labels attached to both S_1_ and R_1_ relative to a positive control of 10 nM S_1_-R_1_-D for a-d and 50 nM S_1_-R_1_-D for e (shown in black dashed line in a-e). All experiments initially have 10 nM concentrations each of S_1_-L, R_1_ and the components of the corresponding upstream logic circuit. Variable concentrations of input and template were then injected, as indicated in the plots, and FRET output reported. (a) YES gate (Figure 5a). (b) AND gate (Figure 5b). (c) YES gate with amplification (Figure 5c). (d) NOT gate implemented via a sequestration of the input to the amplified YES gate in (c) (as sketched in Figure 5d). In this case, the equivalent curves in (c) act as a comparison for the case when the input 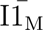 was not added. (e) Amplified YES gate, but using a large ratio of monomers to inputs, each strand of which must then release several fuel molecules on average. (f) Schematics illustrating the placement of fluorophore and quencher labels on the strands and the generation of the FRET signal within the system, which is employed to analyse the reaction mechanism.

For the YES gate in Figure 6a, standard catalytic turnover was observed when both the template and the input I1 were present. In the absence of the template, negligible reaction progress was observed, while in the absence of I1 we observe a sharp initial rise followed by a plateau, corresponding to formation of S_1_-T_11_-R_1_ without the subsequent release of the product, as expected. For the AND gate in Figure 6b, similar results were obtained but now both I1 and I2 are necessary to trigger release of the product and allow catalytic turnover. For the amplified YES gate plotted in Figure 6c, multiple cycles of input I1_M_ - mediated release of D are necessary to drive the reaction to completion, explaining the delayed onset of growth for systems with both I1_M_ and T_11_. In Figure 6d we show that adding 25 nM of the input complement 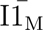 to the amplified YES gate prevents the release of fuel and hence catalytic turnover, as expected. Finally, in Figure 6e, we show that the turnover of the input allows 2nM of input I1_M_ to release enough fuel that ∼ 40 nM of monomers can undergo catalytic templating. Figure 6f shows a schematic representation of strands labelled with fluorophores and quenchers, illustrating the mechanism of FRET signal generation, employed to probe the reaction pathway.

## CONCLUSIONS

We have developed a programmable, enzyme-free system that exhibits sequence-specific catalytic templating of molecular dimerization through DNA strand displacement. To date, there are remarkably few effective mechanisms for catalytic templating of assembly.^5,28^ Here, we demonstrate that the central challenge of product inhibition can be overcome through the use of an ancillary molecule whose involvement is triggered by the bonding of the assembling monomers.

In addition to simply expanding the design space of synthetic templating mechanisms, fuel-driven turnover has some specific advantages. First, the fuel acts as a universal input whose abundance can be used to modulate templating processes, including through upstream molecular networks. Second, it provides more space within which to independently tune the kinetics and thermodynamics of the reaction, which is an essential feature when constructing far-from equilibrium systems such as templating cycles. ^3,35,42^

In our design, the fuel acts through an internal, associative toehold. One might assume that such an involved process might be quite slow, but we observe that a 7 nt toehold is sufficient for reasonable kinetics, at least when assisted by the elimination of mismatches during branch migration. We hypothesise that internal associative toeholds might be useful in other systems, and note while the association that connects the toehold and branch migration in this work is driven by HMSD, alternatives are possible.

Our dimerization systems are still a long way from the power of templating in nature. Sensible next steps would be to look to extend the mechanism to longer templates, and also to use templating to assemble different functional products from a pool labelled monomer units, rather than just products with different size and/or fluorescence properties.

## MATERIALS AND METHODS

### System design

All sequences used are listed in section SI.1 (Supporting Tables S2, S4 and S6). The description of the strands are provided in Supporting Tables S1, S3 and S5. We first designed our DNA reaction networks at the domain level, choosing toeholds of length 6-7 nt, handholds of 7-9 nt, a universal secondary toehold of 1 nt, and internal toeholds ranging from 5 to 10 nt, based on prior work.^5,35^ The branch migration domains were made long enough to contain several mismatches, well separated from each other and well away from duplex ends, to provide thermodynamic drive towards product assembly while minimising unintended leak reaction.^30,43^ Specifically, we incorporated 5 C-C mismatches, which are predicted to be strongly destablising^43^ in the initial blocked S-L monomer at intervals of 6, 7 or 8 base pairs. In the first step (TMSD), when the template-bound monomer 1 is formed, all the five mismatches were retained (Figure 2a). In the second step (HMSD) when the template-bound dimer is formed, three of the mismatches were eliminated (Figure 2b). The other two mismatches were repaired in the third step of the reaction (internal TMSD), when the fuel displaces the template (Figure 2c). Linkers were also included to attach fluorophores to ssDNAs, where appropriate.

Given the above domain architectures and constraints, we obtained corresponding sequences using the Python NUPACK module and NUPACK 4.0 web application.^44,45^ While designing these oligonucleotides, we tried to minimize the secondary structures in the complexes. All strands were purchased from Integrated DNA Technologies (IDT) with high-performance liquid chromatography purification and normalized at 100 µM in LabReady buffer.

### Initial dilution and annealing of complexes

The specific annealing concentrations for each complex were reported in Supporting Tables S9 and S10. Initially we prepared stock solutions of the DNA oligos of ∼ 2 µM concentration in 1X Tris-acetate-EDTA (TAE) buffer from the 100 µM original stock. Absorbance measurements were then performed at wavelength 260 nm, using an IMPLEN N50 nanophotometer, to determine the exact concentrations of the solutions. These recalculated values were further used for annealing or subsequent dilutions.

Strands introduced into experiments directly were prepared at 100 nM using these stock solutions. Working solutions of reactant complexes were prepared by annealing ss- DNAs in a 100 µL volume at a concentration of 200 nM, using the experimental buffer (Tris/acetate/EDTA 1× and 1 M NaCl, pH 8.3). This annealing was performed with an intentional excess of one strand relative to the other, as detailed in Supporting Tables S9 and S10, using a ProFlex PCR System thermocycler by heating to 95 *^◦^*C for 5 min and cooling to 20 *^◦^*C at a rate of 0.5 *^◦^*C min^-1^, and then holding at 4 *^◦^*C until needed. Prior to any experiment, all strand solutions were first incubated at 25 *^◦^*C for at least 20 minutes.

### Fluorescence spectroscopy

Fluorescence assays were carried out in a Clariostar Microplate reader (BMG LABTECH) using Greiner flat µClear bottom 96-well plates, with bottom optics. For the kinetic measurements, we first read the baseline with 150 µL of experimental buffer containing the reactant species, shaken before each cycle for 10 s (double-orbital, 400 rpm). Experimental readings were taken after injecting 50 µL of the triggering reagent at 430 µL/s pump speed and shaking for 10 s after injection. In certain cases, to determine the initial concentration of a reactant species with a quenched fluorophore, we added a large excess of the trigger at the end of the experiment. This excess ensures that 100% of the reactant species is consumed, allowing an estimate of its initial concentration. The reaction protocol for each of the experiments are provided in section SI.2.5 (Supporting Tables S12 - S25).

The reactions were performed at 25 *^◦^*C in 200 µL volume and all the reagents were pre-heated to the experiment temperature. The samples were contained in Eppendorf Lobind tubes, and the injector system of the plate reader was passivated by incubating with 2% (w/v) BSA (bovine serum albumin) in 1X TAE buffer for 15-20 min to maximize concentration reproducibility during the assays.^46^

The kinetic results were obtained by monitoring the fluorescence of fluorophore-labelled strands. The labels used in the experiments were ATTO 488, Alexa Fluor 546, ATTO 425, ATTO 590, and Iowa Black FQ. The plate reader settings for each fluorophore are reported in Supporting Table S11.

The fluorescence signal was averaged for 80 flashes per data point in a spiral area scan with 5 mm radius. We recorded the fluorescence of run-time positive controls and a blank (experimental buffer). These controls were used to correct the measured fluorescence due to physical shaking, temperature changes and volume changes. More detailed protocols for each experiment performed in this work are provided in the Supporting Information (section SI.2). The raw experimental data are available at https://doi.org/10.5281/zenodo.18674501 (see ‘Data availability’ section).

### Gel electrophoresis

As a support for the specificity experiment, we performed gel electrophoresis to assess the orthogonality of templating. For PAGE analysis, the experiments as described in Supporting Table S25 were run for 24 h and each well of the 20% TBE gel was loaded with 10 µL of 10 nM of these reaction samples or controls and 2 µL 6x Purple Loading Dye (no SDS), for a total of 12 µL per lane. This yields a final dye concentration 1x and a sample concentration of 8.3 nM. The controls were pre-annealed complexes. The electrophoresis was performed at a constant 200 V for 2h in 1x TBE buffer. On completion of the run, the gel was washed for 10 min in 1x TBE at 40 rpm to rinse off the excess dye. Then the gel was imaged on an Amersham Typhoon fluorescence gel scanner using the following instrument settings: blue (excitation: 488 nm, emission: 525/20 nm, photomultiplier tube voltage: 344 V), and green (excitation: 532 nm, emission: 570/20 nm, photomultiplier tube voltage: 343 V. The pseudocolour cropped image reported in Figure 4e and the full image reported in Supporting Figure S13 are generated by merging the original greyscale images using ImageJ software, after adjusting the brightness for better visibility and reduced background. The raw fluorescence images of the polyacrylamide gels are available at https://doi.org/10.5281/zenodo.18674501 (see ‘Data availability’ section).

### Data processing

The data from fluorescence experiments were blank corrected and then rescaled by the run-time positive controls for each experiment. They were then transformed from fluorescence units to concentrations of the relevant species where possible using initial data points and signal after saturation, where relevant. For any triplicate data plot, the ‘mean’ represents the arithmetic mean of concentration values at each time point across all replicates in the group and the ‘range’ represents the full range (minimum to maximum) of replicate values at each time point. The transformation procedures are described in detail in section SI.3 and the implementing code Clearissa v.1.71 can be downloaded from https://github.com/PrinciplesKJ/Clearissa_1.71 and the specific instructions are available at https://doi.org/10.5281/zenodo.18674501 (see ‘Data availability’ section). The data processing steps are illustrated in Supporting Figure S1, and the data processed supplementary figures are provided in section SI.5.

### Data fitting

Simple models of reaction kinetics were used to fit reaction rate constants to the processed data in some experiments. All fits are performed with in-house software Clearissa (PYTHON). Supporting Information SI.4 contains a detailed description of the fitting procedures. The fitted data can be generated using the Clearissa software, which is available for download from https://github.com/PrinciplesKJ/Clearissa_1.71. The final figures can be reproduced using the data and instructions uploaded to https://doi.org/10.5281/zenodo.18674501. For operating the Clearissa software (version 1.71), we used Python 3.12.9 on Windows 11 Pro, OS build 26100.7623. On other systems the fitting may vary slightly.

### Data availability

All the experimental data and the code used for processing them are available in the following links.

- Data analysis software: https://github.com/PrinciplesKJ/Clearissa_1.71
- Raw data and processing instructions: https://doi.org/10.5281/zenodo.18674501

## Acknowledgement

M. M., R. M. and T. E. O. are funded by the European Research Council (ERC) under the European Union’s Horizon 2020 research and innovation programme (grant agreement 851910). M. M., K. J. and T. E. O. thank The Royal Society University Research Fellowship Renewal and associated Expenses (grant no. URF\R\211020 to T.E.O., RF\ERE\231045 to M. M. and RF\ERE\210246 to K. J.).

## Supporting Information Available

The Supporting Information is available on ACS Publications website, and provides detailed descriptions of the strands used in the experiments, along with the experimental procedures and reaction protocols. It also includes the methods for data processing and fitting, as well as figures presenting the supplementary results.

## Notes

https://zenodo.org/records/18674501

### Competing Interest Statement

The authors have declared no competing interest.

