## Supplementary notes, figures, and tables for "Fuel-driven catalytic molecular templating"

#### SI.1 DNA strands and sequences

Each strand used in the the reported work is described in Tables S1, S3 and S5. All sequences used are listed in Tables S2, S4 and S6.

##### SI.1.1 Description of DNA sequences for the first catalytic templating system

All strand sequences are written from 5'to 3'. Poly T linkers are used to connect the strands to fluorophores, or to produce different migration rates in PAGE gels, where necessary. Sequences are coloured by domains using the following scheme:

Lilac: Toehold and its complement in monomer 1 and templates.

Red: Branch migration domain and its complement that binds monomer 1 to the fuel.

Black: Branch migration domain and its complement that binds monomer 1 to monomer 2.

Blue: internal toehold in monomer 2 and its complement in the fuel.

Yellow: handold contained in monomer 2 and its complement in the templates.

**Grey and bold: Sequence mismatches whose repair drive the dimerization reaction.**

Table S1. Description of strands used for the first catalytic templating system.

| **Strands** | **Description** | **Variants with labelling** |
|---|---|---|
| $S_1$ | First monomer of the first set. Contains the toehold and is initially blocked. | Two variants:<br><br>a) fluorophore on 3' end;<br>b) fluorophore on 5' end. |
| $R_1$ | Second monomer of the first set. Contains the handhold and internal toehold. | Two variants:<br><br>a) fluorophore on 5' end;<br>b) unlabelled. |
| L | Blocker that initially blocks monomer $S_1$. | Quencher on 5' end. |
| $T_{11}$ (also called $T^{8\text{-}7}$) | Template $T_{11}$ with handhold length 8 nt and toehold length 7 nt and dimerizes $S_1$ and $R_1$. | Two variants:<br><br>a) unlabelled;<br>b) quencher on 3' end. |
| $T^{p\text{-}q}$ | To test variants of toehold and handhold lengths, we used templates $T^{p\text{-}q}$, where 'p' is the length of the handhold, and 'q' is the length of toehold. | Unlabelled. |
| $D^j$ | Fuel with internal toehold length 'j'. | Unlabelled. |

Table S2. DNA sequences used for the first catalytic templating system.

| **Strands** | **DNA Sequences** |
|---|---|
| $S_1$(a) | CCCTTAACCTACCCTCACATCTCCACCTCAACCTAACTCCTAACTACTACTGGCC TTT ATTO 488 |
| $S_1$(b) | ATTO 488 TTT CCCTTAACCTACCCTCACATCTCCACCTCAACCTAACTCCTAACTACTACTGGCC |
| $R_1$(a) | Alexa 546 TTT GGCCAGTAGTAGTTAGGAGTTAGGTCCATCATCCATCAAATTCAC |
| $R_1$(b) | GGCCAGTAGTAGTTAGGAGTTAGGTCCATCATCCATCAAATTCAC |
| L | Iowa Black FQ TTT GGCCAGTACTAGTTACGAGTTACGTTAGGTGCAGATGTCAGGGTAGG |
| $T_{11}/T^{8\text{-}7}$(a) | GAATTTGAGCCAGTACTAGTTACGAGTTACGTTAGGTGCAGATGTCAGGGTAGGTTAAGGG |
| $T_{11}/T^{8\text{-}7}$(b) | GAATTTGAGCCAGTACTAGTTACGAGTTACGTTAGGTGCAGATGTCAGGGTAGGTTAAGGG TTT Iowa Black FQ |
| $T^{9\text{-}7}$ | TGAATTTGAGCCAGTACTAGTTACGAGTTACGTTAGGTGCAGATGTCAGGGTAGGTTAAGGG |
| $T^{9\text{-}6}$ | TGAATTTGAGCCAGTACTAGTTACGAGTTACGTTAGGTGCAGATGTCAGGGTAGGTTAAGG |
| $T^{8\text{-}6}$ | GAATTTGAGCCAGTACTAGTTACGAGTTACGTTAGGTGCAGATGTCAGGGTAGGTTAAGG |
| $T^{7\text{-}7}$ | AATTTGAGCCAGTACTAGTTACGAGTTACGTTAGGTGCAGATGTCAGGGTAGGTTAAGGG |
| $T^{7\text{-}6}$ | AATTTGAGCCAGTACTAGTTACGAGTTACGTTAGGTGCAGATGTCAGGGTAGGTTAAGG |
| $T^{0\text{-}7}$ | GCCAGTACTAGTTACGAGTTACGTTAGGTGCAGATGTCAGGGTAGGTTAAGGG |
| $D^{10}$ | TGGATGATGGTAGGTGGAGATGTGAGGGTAGG |
| $D^7$ | ATGATGGTAGGTGGAGATGTGAGGGTAGG |
| $D^6$ | TGATGGTAGGTGGAGATGTGAGGGTAGG |
| $D^5$ | GATGGTAGGTGGAGATGTGAGGGTAGG |

##### SI.1.2 DNA strands used to demonstrate orthogonality in the catalytic templating system

All the templates used in these reactions have handhold of length 8 nt and toehold 7 nt. These strands were used along with $S_1$, $R_1$, L, $T_{11}$, $D^7$, as described in Table S1. Sequences are coloured using the same colour scheme.

Table S3. Strands used to demonstrate orthogonality in the catalytic templating system.

| Strands | Description | Variants with labelling |
|---|---|---|
| $S_2$ | First monomer of the second. Contains a toehold orthogonal to that of $S_1$, but the same branch migration domains. | Two variants: a) fluorophore on 3' end; b) fluorophore on 5' end. |
| $R_2$ | Second monomer of the second set. Contains a handhold orthogonal to that of $R_1$, but the same internal toehold and branch migration domain. | Unlabelled |
| $T_{22}$ | Template $T_{22}$ that dimerizes $S_2$ and $R_2$ | Two variants: a) unlabelled; b) quencher on 3' end. |
| $T_{12}$ | Template $T_{12}$ that dimerizes $S_1$ and $R_2$ | Two variants: a) unlabelled; b) quencher on 3' end. |
| $T_{21}$ | Template $T_{21}$ that dimerizes $S_2$ and $R_1$ | Two variants: a) unlabelled; b) quencher on 3' end. |

Table S4. DNA sequences to demonstrate orthogonal catalytic templating.

| Strands | DNA Sequences |
|---|---|
| $S_2$(a) | CGCTCTACCTACCCTCACATCTCCACCTCAACCTAACTCCTAACTACTACTGGCC TTT Alexa 546 |
| $S_2$(b) | Alexa 546 TTT CGCTCTACCTACCCTCACATCTCCACCTCAACCTAACTCCTAACTACTACTGGCC |
| $R_2$ | TTT GGCCAGTAGTAGTTAGGAGTTAGGTCCATCATCCATCCATCTCAA |
| $T_{22}$(a) | GAGATGGAGCCAGTACTAGTTACGAGTTACGTTAGGTGCAGATGTCAGGGTAGGTAGAGCG |
| $T_{22}$(b) | GAGATGGAGCCAGTACTAGTTACGAGTTACGTTAGGTGCAGATGTCAGGGTAGGTAGAGCG TTT Iowa Black FQ |
| $T_{12}$(a) | GAGATGGAGCCAGTACTAGTTACGAGTTACGTTAGGTGCAGATGTCAGGGTAGGTTAAGGG |
| $T_{12}$(b) | GAGATGGAGCCAGTACTAGTTACGAGTTACGTTAGGTGCAGATGTCAGGGTAGGTTAAGGG TTT Iowa Black FQ |
| $T_{21}$(a) | GAATTTGAGCCAGTACTAGTTACGAGTTACGTTAGGTGCAGATGTCAGGGTAGGTAGAGCG |
| $T_{21}$(b) | GAATTTGAGCCAGTACTAGTTACGAGTTACGTTAGGTGCAGATGTCAGGGTAGGTAGAGCG TTT Iowa Black FQ |

##### SI.1.3 Description of strands used to demonstrate upstream control of catalytic templating

The strands described in Table S5 were used along with $S_1$(a), $R_1$(a), L and $T_{11}$(a), as described in Table S1.

Sequences in Table S6 are coloured by domain using the following colours:

Teal: extension of fuel Dn that allows it to bind with with X or XE, sand its complement. Branch migration domain of inputs I1, $I1_M$, Z and $I\bar{1}_M$.

Blue: internal toehold in fuel Dn.

Red: Branch migration domain of fuel Dn that allows it to bind to $S_1$.

Orange: toehold and its complement in X, XE, and inputs I1, $I1_M$, $I\bar{1}_M$.

Lavender: branch migration domain and its complement in input I2 and XE.

Navy: toehold and its complement in Input I2 and XE

**Grey and bold: mismatches**

Table S5. Description of strands used for the upstream gate circuitry.

| **Strands** | **Description** | **Used in gated systems** | **Variants with labelling** |
|---|---|---|---|
| Dn | An extended fuel strand with internal toehold length 7 nt which is initially bound to X or XE | all | unlabelled |
| X | Fuel Dn is bound to strand X | Single-input, catalytic, NOT | quencher on 5' end |
| I1 | Input strand | Single-input, 2-input AND | fluorophore on 3' end |
| E | Initially bound to XE strand, which detaches to open up a toehold for the input I2 | 2-input AND | fluorophore on 5' end |
| XE | Fuel Dn and strand E is initially bound to XE | 2-input AND | quencher on 5' end |
| I2 | Input strand | 2-input AND | unlabelled |
| $\mathrm{I1}_{\mathrm{M}}$ | Input strand, which is similar to input I1, but with one nt change (C instead of G at 24th position) to retain a mismatch in the second step of the reaction | catalytic, NOT | fluorophore on 3' end |
| Z | Ancillary strand | catalytic, NOT | unlabelled |
| $\mathrm{I}\bar{1}_{\mathrm{M}}$ | Input strand – fully complementary to $\mathrm{I1}_{\mathrm{M}}$ | NOT | unlabelled |

Table S6. DNA sequences used for the upstream gate circuitry.

| **Strands** | **DNA Sequences** |
|---|---|
| Dn | GGCGATTTCAGTGGAGCTTTAGTTGATGATGGTAGGTGGAGATGTGAGGGTAGG |
| X | Iowa Black FQ TTT CCATCATCAACTAAACCTCCACTCAAATCGCCAAATAGA |
| I1 | TCTATTTGGCGATTTGAGTGGAGGTTTAGTTG TTT ATTO 590 |
| E | ATTO 425 TTT GTAGTTATCATTTGGCCAGGTTAGTTCTATTT |
| XE | Iowa Black FQ TTT<br>CCATCATCAACTAAACCTCCACTCAAATCGCCAAATAGAACTAACCTCGCCAAATCATAACTACACTCACC |
| I2 | GGTGAGTGTAGTTATGATTTGGCGAGGTTAGT |
| $\mathrm{I1}_{\mathrm{M}}$ | TCTATTTGGCGATTTGAGTGGAGCTTTAGTTG TTT ATTO 590 |
| Z | GGCGATTTGAGTGGAGGTTTAGTTGATGATGG |
| $\mathrm{I}\bar{1}_{\mathrm{M}}$ | CAACTAAAGCTCCACTCAAATCGCCAAATAGA |

##### SI.1.4 DNA strands used for experiments shown in figures

The strands used in the reported figures of both the main text and the supplementary information are listed in Tables S7 and S8.

Table S7. DNA strands used in experiments reported in the main text.

| **Figure numbers** | **Strands used** |
|---|---|
| 2a | $S_1$(a), L, $T_{11}$(a) |
| 2b | $S_1$(a), $T_{11}$(a), $R_1$(a) |
| 2c | $S_1$(b), $T_{11}$(b), $R_1$(b), $D^7$ |
| 3a | $S_1$(a), L, $R_1$(b), $D^5$, $T_{11}$(a) |
| 3b | $S_1$(a), L, $R_1$(b), $D^6$, $T_{11}$(a) |
| 3c | $S_1$(a), L, $R_1$(b), $D^7$, $T_{11}$(a) |
| 3d | $S_1$(a), L, $R_1$(b), $D^{10}$, $T_{11}$(a) |
| 3e | $S_1$(a), L, $R_1$(b), $D^6$, $T_{11}$(a) |
| 3f | $S_1$(a), L, $R_1$(b), $D^7$, $T_{11}$(a) |
| 4a | $S_1$(a), L, $R_1$(b), $D^7$, $T_{11}$(a), $T_{12}$(a), $T_{21}$(a), $T_{22}$(a) |
| 4b | $S_1$(a), L, $R_1$(b), $S_2$(a), $R_2$, $D^7$, $T_{11}$(a), $T_{12}$(a), $T_{21}$(a), $T_{22}$(a) |
| 4c | $S_1$(a), L, $R_1$(b), $S_2$(a), $R_2$, $D^7$, $T_{11}$(a), $T_{12}$(a), $T_{21}$(a), $T_{22}$(a) |
| 4e | $S_1$(a), $S_2$(a), L, $R_1$(b), $R_2$, $D^7$, $T_{11}$(a), $T_{12}$(a), $T_{21}$(a), $T_{22}$(a) |
| 6a | Dn, X, I1, $S_1$(a), L, $R_1$(a), $T_{11}$(a) |
| 6b | Dn, E, XE, I2, I1, $S_1$(a), L, $R_1$(a), $T_{11}$(a) |
| 6c | Dn, X, $I1_M$, Z, $S_1$(a), L, $R_1$(a), $T_{11}$(a) |
| 6d | Dn, X, $I1_M$, Z, $I\bar{1}_M$, $S_1$(a), L, $R_1$(a), $T_{11}$(a) |
| 6e | Dn, X, $I1_M$, Z, $S_1$(a), L, $R_1$(a), $T_{11}$(a) |

Table S8. DNA strands used in experiments reported in supporting figures.

| **Figure numbers** | **Strands used** |
|---|---|
| S2a | $S_1$(a), L, $T^{8\text{-}6}$ |
| S2b | $S_1$(a), L, $T^{8\text{-}7}$ |
| S3a | $S_1$(a), $T^{0\text{-}7}$, $R_1$(a) |
| S3b | $S_1$(a), $T^{7\text{-}7}$, $R_1$(a) |

| | |
|---|---|
| S3c | $S_1$(a), $T^{8\text{-}7}$(a), $R_1$(a) |
| S3d | $S_1$(a), $T^{9\text{-}7}$, $R_1$(a) |
| S4a | $S_1$(b), $T_{11}$(b), $R_1$(b), $D^5$ |
| S4b | $S_1$(b), $T_{11}$(b), $R_1$(b), $D^6$ |
| S4c | $S_1$(b), $T_{11}$(b), $R_1$(b), $D^{10}$ |
| S5 | $S_1$(a), $T_{11}$(a), L |
| S6a | $S_1$(b), $R_1$(b), $D^5$, $T_{11}$(b) |
| S6b | $S_1$(b), $R_1$(b), $D^6$, $T_{11}$(b) |
| S6c | $S_1$(b), $R_1$(b), $D^7$, $T_{11}$(b) |
| S6d | $S_1$(b), $R_1$(b), $D^{10}$, $T_{11}$(b) |
| S7a | $S_1$(b), $R_1$(b), $D^7$, $T_{11}$(b) |
| S7b | $S_1$(b), $R_2$, $D^7$, $T_{12}$(b) |
| S7c | $S_2$(b), $R_1$(b), $D^7$, $T_{21}$(b) |
| S7d | $S_2$(b), $R_2$, $D^7$, $T_{22}$(b) |
| S8a | $S_1$(a), L, $R_1$(a), $D^7$, $T^{9\text{-}7}$ |
| S8b | $S_1$(a), L, $R_1$(a), $D^7$, $T^{8\text{-}7}$ |
| S8c | $S_1$(a), L, $R_1$(a), $D^7$, $T^{7\text{-}7}$ |
| S8d | $S_1$(a), L, $R_1$(a), $D^7$, $T^{9\text{-}6}$ |
| S8e | $S_1$(a), L, $R_1$(a), $D^7$, $T^{8\text{-}6}$ |
| S8f | $S_1$(a), L, $R_1$(a), $D^7$, $T^{7\text{-}6}$ |
| S9a | $S_1$(a), L, $R_1$(b), $D^7$, $T_{11}$(a) |
| S9b | $S_1$(a), L, $R_1$(b), $D^6$, $T_{11}$(a) |
| S10a-b | $S_1$(a), L, $R_1$(b), $D^7$, $T_{11}$(a) |
| S11 | $S_2$(a), L, $R_2$, $D^7$, $T_{22}$(a) |
| S12a | $S_1$(a), L, $R_2$, $D^7$, $T_{11}$(a), $T_{12}$(a), $T_{21}$(a), $T_{22}$(a) |
| S12b | $S_2$(a), L, $R_1$(b), $D^7$, $T_{11}$(a), $T_{12}$(a), $T_{21}$(a), $T_{22}$(a) |
| S12c | $S_2$(a), L, $R_2$, $D^7$, $T_{11}$(a), $T_{12}$(a), $T_{21}$(a), $T_{22}$(a) |
| S13 | $S_1$(a), $S_2$(a), L, $R_1$(b), $R_2$, $D^7$, $T_{11}$(a), $T_{12}$(a), $T_{21}$(a), $T_{22}$(a) |
| S14 | $R_1$(a), $D^{10}$, $D^7$, $D^6$ |

### SI.2 Experimental procedures

#### SI.2.1 Annealing of complexes

Annealing was performed according to the protocol reported in the methods section of the manuscript. Working solutions were prepared with a final concentration as tabulated in Tables S9 – S10. Excesses of one strand were used to ensure all complementary partners were bound.

Table S9. Concentrations of DNA strands in working solutions of annealed complexes used to demonstrate catalytic templating in both the orthogonal sets.

| **Complexes** | **Strand names** | **Strand annotations** | **Final conc (nM)** | **Description** |
|---|---|---|---|---|
| Reactant $S_1$-L/$S_2$-L | Monomer 1 | $S_1$(a)/$S_2$(a) | 200 | |
| | Blocker | L | 400 | 100% excess |
| Control $S_1$-T | Monomer 1 | $S_1$(a) | 200 | |
| | Template | $T_{11}$(a)/$T^{p\text{-}q}$ | 240 | 20% excess |
| Reactant $S_1$-T | Monomer 1 | $S_1$(a) | 200 | |
| | Template | $T_{11}$(a)/$T^{p\text{-}q}$ | 300 | 50% excess |
| Control $S_1$-$R_1$-T | Monomer 1 | $S_1$(a) | 200 | |
| | Monomer 2 | $R_1$(a) | 240 | 20% excess |
| | Template | $T_{11}$(a)/$T^{p\text{-}q}$ | 240 | 20% excess |
| Reactant $S_1$-$R_1$-T | Monomer 1 | $S_1$(b) | 200 | |
| | Monomer 2 | $R_1$(b) | 300 | 50% excess |
| | Template | $T_{11}$(b) | 300 | 50% excess |
| Controls $S_1$-$R_1$-D, $S_1$-$R_2$-D, $S_2$-$R_1$-D, $S_2$-$R_2$-D | Monomer 1 | $S_1$(a)/$S_1$(b)/ $S_2$(a)/$S_2$(b) | 200 | |
| | Monomer 2 | $R_1$(a)/$R_1$(b)/$R_2$ | 240 | 20% excess |
| | Fuel | $D^j$ | 240 | 20% excess |

Table S10. Concentrations of DNA strands in working solutions of annealed complexes used to demonstrate catalytic templating coupled with the upstream DNA gate circuitry.

| **Strand names** | **Strand annotations** | **Final conc (nM)** | **Description** |
|---|---|---|---|
| Dn-X | Dn | 200 | |
| | X | 220 | 10% excess |
| E-Dn-XE | Dn | 200 | |
| | E | 240 | 20% excess |
| | XE | 220 | 10% excess |
| Control $S_1$-$R_1$-Dn | $S_1$(a) | 200 | |
| | $R_1$(a) | 240 | 20% excess |
| | Dn | 240 | 20% excess |

#### SI.2.2 Controls

In all experiments, we used four different concentrations for the positive control (run-time control), except the specificity experiments shown in Table S23 and S24 where we have used two different concentrations for each set of reactions; at least one negative control (where the triggering agent/(s) is/are absent) and a blank consisting of 1x TAE buffer (working buffer). The controls were subjected to identical volume changes and environmental conditions as those experienced by the samples during the kinetic measurements. These controls enabled correction for fluorescence changes arising from environmental variations, and the procedure for data correction using the control signals is described in section SI.3.2.

##### SI.2.3 Preparation of plate and reading with the plate reader

At the start of the experiment, all the reactants (except the triggering agent/s) were added at the desired concentrations and volumes to the experimental buffer (1M NaCl in 1x TAE) to obtain a total reaction volume of 150 µL. The experimental concentrations reported in both the main text and supplementary information represent the final concentrations in a total volume of 200 µL, accounting for the additional 50 µL that was introduced during the injection step. All the different fluorescence measurements in the different experiments followed a similar structure, outlined below:

**Measurement 1.** Initial fluorescence: We took an initial measurement (baseline) with incubation at 25 °C to gather at least 20 data points after the fluorescence signals of the positive controls became stable. This approach allows us to minimize the uncertainties in concentration that may result from factors such as temperature variations, pipetting errors, and other experimental inconsistencies.

**Measurement 2.** Injection of triggering agent and kinetic measurement: For the majority of experiments, 50 µL of the triggering reactant was injected by the plate reader, bringing the total reaction volume to 200 µL. For all but the fastest reactions, the plate was then completely sealed to prevent evaporation, after a short period in which the initial dynamics was monitored carefully. The fluorescence signal was then monitored for long enough to reach steady state for all but the slowest of reactions. In some experiments with slower kinetics in which multiple different triggers were added to the same plate, the triggering was instead performed by hand, the plate sealed, and added to the plate reader. The endpoint fluorescence of measurement 2 can be used to infer concentrations of certain initial reactants (assuming irreversibility and reaction completion).

**Measurement 3.** Saturation: To finish the assay, experiments were often saturated with an excess (which must not be any fluorophore-containing species) of a non-fluorescent reagent. This saturation allowed an estimate of the initial concentration of reactants where this information is required.

All the procedures mentioned above are illustrated in Figure S1 and the measurements are annotated as m1-m3.

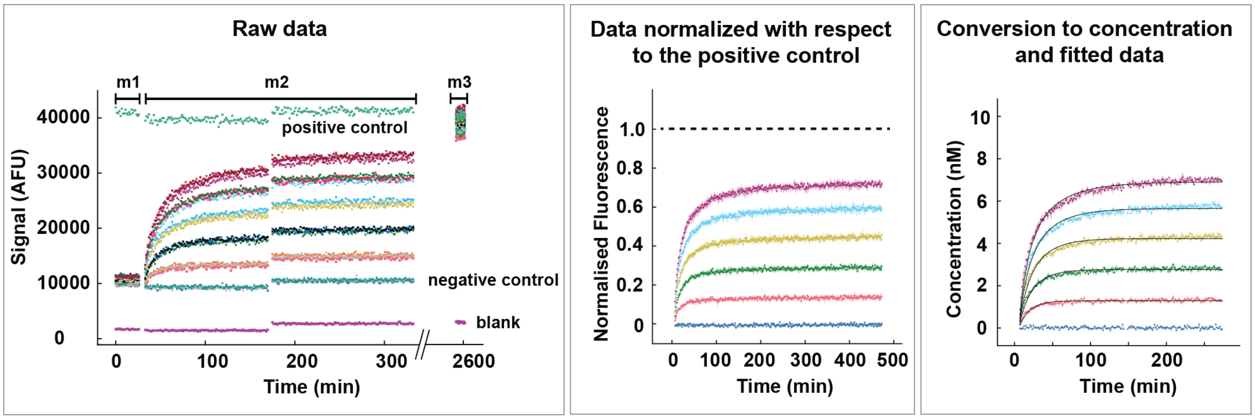

Figure S1. Demonstration of a complete set of measurements for a general reaction using a Microplate reader and the processing of data. In the left panel, 'm1' represents the baseline of the experiments, considered as 0 nM of reacted species, before the injection of the triggering agent. At the start of 'm2', varying amounts of the triggering reactant were added and the reaction was then monitored. The plate was sealed at around 175 minutes, and then monitored for a further few hours to complete m2. Afterwards, 'm3' was taken where the reaction mixtures were saturated with a large excess of the non-fluorescent reactant, sufficient enough so that the fluorophore-labelled species react completely. The negative controls consists of all the reactants, except the triggering agent. Although 4 postive controls at different concentrations were used in experiments, we plot only one here for ease of interpretation. After data processing (section SI.3) and kinetic fitting (section SI.4) the figures (middle and right) in the subsequent panels are obtained.

##### SI.2.4 Fluorescence measurement settings

The kinetic measurements were obtained by monitoring the fluorescence of fluorophore-labelled strands. The experimental settings for each fluorophore are reported in Table S11.

Table S11. Settings for monitoring of fluorescence

| **Fluorophores** | **Excitation** | **Dichroic filter** | **Emission** | **Gain** | **Gain for high conc samples** |
|---|---|---|---|---|---|
| ATTO 488 | 488–14 | 504.8 | 519–9 | 2500 | 2200 |
| Alexa Fluor 546 | 540–20 | 562.5 | 590–30 | 1800 | 1400 |
| FRET: ATTO 488 – Alexa Fluor 546 | 488–10 | 534 | 590–30 | 2000 | 1600 |
| ATTO 425 | 430–15 | 461.2 | 490–40 | 2400 | – |
| ATTO 590 | 575–30 | 602 | 634–40 | 2000 | – |

The specific procedures for each type of experiment are discussed and collected in Tables in the following subsections.

#### SI.2.5 Protocols for each experiment

In Tables S12 - S30, each **concentration** column represents a single reaction well, containing the reactants – listed row by row – in the intended concentrations in 200 µL volume.

**A. Step 1 – toehold-mediated strand displacement.** Increasing concentrations (0–10 nM) of the template $T^{p\text{-}q}$ were added to wells containing 10 nM of pre-annealed, blocked monomer $S_1$(a)-L. After the signal reached a plateau, all reaction wells were treated with large excess of $T^{p\text{-}q}$ to ensure complete consumption of any unreacted monomer in the reaction mixture. The reaction conditions are tabulated in Table S12.

Table S12. Reaction protocol for step 1 for experiments shown in Figures 2a in the manuscript and S2.

| Reactants | Concentrations in nM | | | | | |
|---|---|---|---|---|---|---|
| $S_1$-L | 10 | 10 | 10 | 10 | 10 | 10 |
| $T^{p\text{-}q}$ (added by injector) | 0 | 2 | 4 | 6 | 8 | 10 |

| Positive Control | Concentrations in nM | | | |
|---|---|---|---|---|
| $S_1$-$T^{p\text{-}q}$ | 5 | 10 | 15 | 20 |

**B. Step 2 – handhold-mediated strand displacement.** To investigate the second step, monomer $S_1(a)$ and template $T^{p\text{-}q}(a)$ were pre-annealed. Variable concentrations (0–10 nM) of $R_1(a)$ were added to wells containing 10 nM of pre-annealed, template bound monomer $S_1(a)$. The reaction conditions are tabulated in Table S13.

Table S13. Reaction protocol for step 2 for experiments shown in Figures 2b in the manuscript and S3.

| **Reactants** | **Concentrations in nM** | | | | | |
|---|---|---|---|---|---|---|
| $S_1$-$T^{p\text{-}q}$ | 10 | 10 | 10 | 10 | 10 | 10 |
| $R_1$ (added by injector) | 0 | 2 | 4 | 6 | 8 | 10 |

| **Positive Control** | **Concentrations in nM** | | | |
|---|---|---|---|---|
| $S_1$-$R_1$-$T^{p\text{-}q}$ | 5 | 10 | 15 | 20 |

**C. Step 3 – internal toehold-mediated strand displacement.** 10 nM of the pre-annealed complex $S_1(b)$-$R_1(b)$-$T_{11}(b)$ were added to the varying concentrations (0–10 nM) of the fuels $D^j$. After the signal reached a plateau, we saturated the reaction mixtures with excess of $D^j$ in respective wells to ensure complete consumption of any unreacted complex $S_1$-$R_1$-$T_{11}$ in the reaction mixture. The reaction conditions are tabulated in Table S14.

Table S14. Reaction protocol for step 3 for experiments shown in Figures 2c in the manuscript and S4.

| **Reactants** | **Concentrations in nM** | | | | | |
|---|---|---|---|---|---|---|
| $D^j$ | 0 | 2 | 4 | 6 | 8 | 10 |
| $S_1$-$R_1$-$T_{11}$ (added by injector) | 10 | 10 | 10 | 10 | 10 | 10 |

| **Positive Control** | **Concentrations in nM** | | | |
|---|---|---|---|---|
| $S_1$-$R_1$-$D^j$ | 5 | 10 | 15 | 20 |

###### SI.2.5.1 Test for reversibility

To check the thermodynamic drive associated with the templating reaction, we examined the irreversibility of product formation for step 1 and step 3 in the catalytic cycle.

**A. Reversibility of step 1.** We reacted 10 nM of $S_1$(a)-$T_{11}$(a) with variable concentrations (0-10 nM) of the blocker L. The reaction conditions are tabulated in Table S15.

Table S15. Reaction protocol for the reversibility test for step 1 for experiments shown in Figure S5.

| **Reactants** | **Concentrations in nM** | | | | | |
|---|---|---|---|---|---|---|
| $S_1$-$T_{11}$ | 10 | 10 | 10 | 10 | 10 | 10 |
| L (added by injector) | 0 | 2 | 4 | 6 | 8 | 10 |

| **Positive Control** | **Concentrations in nM** | | | |
|---|---|---|---|---|
| $S_1$-L | 5 | 10 | 15 | 20 |

**B. Reversibility of step 3.** We first reacted 10 nM of pre-annealed complex $S_1$(b)-$R_1$(b)-$D^j$ (j = 5, 6, 7 and 10) with variable concentrations (0-10 nM) of $T_{11}$(b). We then repeated the exercise using all 4 orthogonal templates and their target products for $D^7$. For these reactions, pre-annealed complexes $S_1$(b)-$R_1$(b)-D, $S_1$(b)-$R_2$-D, $S_2$(b)-$R_1$(b)-D and $S_2$(b)-$R_2$-D were individually reacted to $T_{11}$(b), $T_{12}$(b), $T_{21}$(b) and $T_{22}$(b) respectively. The reaction conditions are tabulated in Table S16.

Table S16. Reaction protocol for the reversibility test for step 3 for experiments shown in Figures S6 and S7.

| **Reactants** | **Concentrations in nM** | | | | | |
|---|---|---|---|---|---|---|
| $S_k$-$R_l$-$D^j$ | 10 | 10 | 10 | 10 | 10 | 10 |
| $T_{kl}$ (added by injector) | 0 | 2 | 4 | 6 | 8 | 10 |

| **Positive Control** | **Concentrations in nM** | | | |
|---|---|---|---|---|
| $S_k$-$R_l$-$T_{kl}$ | 5 | 10 | 15 | 20 |

##### SI.2.5.2 Optimization of catalytic templating turnover

**A. Optimization of the toehold and the handhold lengths.** Varying concentrations (0–5 nM) of the template $T^{p\text{-}q}$ were added to wells containing 10 nM of each of pre-annealed complex $S_1$(a)-L, monomer $R_1$(a), and fuel $D^7$, considering 6-7 for q and 7-9 for p. The reaction procedure is tabulated in Table S17.

Table S17. Reaction protocol for catalytic templating with templates of varying toehold and handhold lengths for experiments shown in Figure S8.

| Reactants | Concentrations in nM | | | | |
|---|---|---|---|---|---|
| $S_1$-L | 10 | 10 | 10 | 10 | 10 |
| $R_1$ | 10 | 10 | 10 | 10 | 10 |
| $D^7$ | 10 | 10 | 10 | 10 | 10 |
| $T^{p\text{-}q}$ (added by injector) | 0 | 2 | 3 | 4 | 5 |

| Positive Control | Concentrations in nM | | | |
|---|---|---|---|---|
| $S_1$-$R_1$-$D^7$ | 5 | 10 | 15 | 20 |

**B. Optimization of the internal toehold length.** We added varying concentrations of template $T^{8\text{-}7}$ (also known as $T_{11}$) to solutions containing 10 nM each of annealed $S_1$(a)-L, $R_1$(b) and $D^j$, with j = 5, 6, 7 and 10. After the signal reached a plateau, all reaction wells were treated with large excess of $T_{11}$ to ensure complete consumption of any unreacted monomer in the reaction mixture. The concentrations of the reactants are as given in Table S18.

Table S18. Reaction protocol for catalytic templating with fuels of varying internal toehold lengths for experiments shown in Figure 3a-d in the manuscript.

| **Reactants** | **Concentrations in nM** | | | | |
|---|---|---|---|---|---|
| $S_1$-L | 10 | 10 | 10 | 10 | 10 |
| $R_1$ | 10 | 10 | 10 | 10 | 10 |
| $D^j$ | 10 | 10 | 10 | 10 | 10 |
| $T_{11}$ (added by injector) | 0 | 2 | 3 | 4 | 5 |

| **Positive Control** | **Concentrations in nM** | | | |
|---|---|---|---|---|
| $S_1$-$R_1$-$D^7$ | 5 | 10 | 15 | 20 |

###### SI.2.5.3 Catalytic templating regulated by excess of fuel molecule

Pre-annealed $S_1$(a)-L, $R_1$(b) and 2 nM of $T_{11}$ were reacted with excess of fuel $D^7$ or $D^6$ as described in Table S19. To probe the region between 10 nM and 50 nM for $D^6$ further, additional experiments were performed as described in Table S20. In this experiment, after the signal reached a plateau, all reaction wells were treated with large excess of $T_{11}$ to ensure complete consumption of any unreacted monomer in the reaction mixture.

Table S19. Reaction protocol for control of catalytic turnover with excess concentration of fuel for experiments shown in Figure S9.

| **Reactants** | **Concentrations in nM** | | | | | |
|---|---|---|---|---|---|---|
| $S_1$-L | 10 | 10 | 10 | 10 | 10 | 10 |
| $R_1$ | 10 | 10 | 10 | 10 | 10 | 10 |
| $D^7/D^6$ | 0 | 5 | 10 | 50 | 100 | 200 |
| $T_{11}$ (added by injector) | 2 | 2 | 2 | 2 | 2 | 2 |

| **Positive Controls** | **Concentrations in nM** | | | |
|---|---|---|---|---|
| $S_1$-$R_1$-$D^7$ | 5 | 10 | 15 | 20 |
| $S_1$-$R_1$-$D^6$ | 5 | 10 | 15 | 20 |

Table S20. Reaction protocol for control of catalytic turnover with excess concentration of $D_6$ for experiments shown in Figure 3e in the manuscript.

| **Reactants** | **Concentrations in nM** | | | | | |
|---|---|---|---|---|---|---|
| $S_1$-L | 10 | 10 | 10 | 10 | 10 | 10 |
| $R_1$ | 10 | 10 | 10 | 10 | 10 | 10 |
| $D^6$ | 0 | 10 | 20 | 30 | 40 | 50 |
| $T_{11}$ (added by injector) | 2 | 2 | 2 | 2 | 2 | 2 |

| **Positive Control** | **Concentrations in nM** | | | |
|---|---|---|---|---|
| $S_1$-$R_1$-$D^6$ | 5 | 10 | 15 | 20 |

##### SI.2.5.4 Catalytic templating with high monomer-to-template ratio

Pre-annealed $S_1$(a)-L, $R_1$(b) and fuel D[7] were reacted with increasing concentrations of the template $T_{11}$(a). The concentrations of the reactants are given in the table S21.

Table S21. Reaction protocol for experiments demonstrating catalytic turnover with high monomer to template ratio for experiments shown in Figure 3f in the manuscript and S10.

| **Reactants** | **Concentrations in nM** | | | | |
|---|---|---|---|---|---|
| $S_1$-L | 50 | 50 | 50 | 50 | 50 |
| $R_1$ | 50 | 50 | 50 | 50 | 50 |
| D | 100 | 100 | 100 | 100 | 100 |
| $T_{11}$ (added by injector) | 0 | 0.5 | 1.0 | 1.5 | 2.0 |

| **Positive Control** | **Concentrations in nM** | | | |
|---|---|---|---|---|
| $S_1$-$R_1$-D | 10 | 20 | 50 | 100 |

##### SI.2.5.5 Reactions for orthogonal dimerization

**A. Full reaction for orthogonal set 2.** Varying concentrations (0-5 nM) of $T_{22}$ were added to 10 nM of each of pre-annealed complex $S_2$(a)-L, monomer $R_2$ and the fuel $D^7$. The concentrations of the reactants are tabulated in table S22.

Table S22. Reaction protocol for the catalytic templating of the orthogonal set 2 for experiments shown in Figure S11.

| **Reactants** | | | **Concentrations in nM** | | |
|---|---|---|---|---|---|
| $S_2$-L | 10 | 10 | 10 | 10 | 10 |
| $R_2$ | 10 | 10 | 10 | 10 | 10 |
| D | 10 | 10 | 10 | 10 | 10 |
| $T_{22}$ (added by injector) | 0 | 2 | 3 | 4 | 5 |

| **Positive Control** | | **Concentrations in nM** | | |
|---|---|---|---|---|
| $S_2$-$R_2$-D | 5 | 10 | 15 | 20 |

**B. Dimerization reactions for each pair of monomers triggered with four different templates** We first tested specificity by exposing all 4 possible combinations of monomers to each template individually. Pre-annealed $S_1$(a)-L/$S_2$(a)-L, $R_1$(b)/$R_2$ and fuel D were reacted with low concentrations of each template in separate experiments, as detailed in Table S23.

Table S23. Reaction protocol for testing specificity by adding four different templates to each monomer pair for experiments shown in Figure 4a in the manuscript (with $S_1$-$R_1$-D) and Figure S12 (with $S_1$-$R_2$-D, $S_2$-$R_1$-D and $S_2$-$R_2$-D).

| **Reactants** | **Concentrations in nM** | | | |
|---|---|---|---|---|
| $S_1$-L | 10 | 10 | 10 | 10 |
| $R_1$ | 10 | 10 | 10 | 10 |
| D | 10 | 10 | 10 | 10 |
| Template (added manually) | 2 ($T_{11}$) | 2 ($T_{12}$) | 2 ($T_{21}$) | 2 ($T_{22}$) |
| Matching | *th, hh* | *th* | *hh* | – |
| Expected nature of reaction | catalytic turnover | only TMSD | no reaction | no reaction |

| **Positive Control** | | **Concentrations in nM** |
|---|---|---|
| $S_1$-$R_1$-D | 5 | 10 |

| **Reactants** | **Concentrations in nM** | | | |
|---|---|---|---|---|
| $S_1$-L | 10 | 10 | 10 | 10 |
| $R_2$ | 10 | 10 | 10 | 10 |
| D | 10 | 10 | 10 | 10 |
| Template (added manually) | 2 ($T_{11}$) | 2 ($T_{12}$) | 2 ($T_{21}$) | 2 ($T_{22}$) |
| Matching | *th* | *th, hh* | – | *hh* |
| Expected nature of reaction | only TMSD | catalytic turnover | no reaction | no reaction |

| **Positive Control** | | **Concentrations in nM** |
|---|---|---|
| $S_1$-$R_2$-D | 5 | 10 |

| Reactants | Concentrations in nM | | | |
|---|---|---|---|---|
| $S_2$-L | 10 | 10 | 10 | 10 |
| $R_1$ | 10 | 10 | 10 | 10 |
| D | 10 | 10 | 10 | 10 |
| Template (added manually) | 2 ($T_{11}$) | 2 ($T_{12}$) | 2 ($T_{21}$) | 2 ($T_{22}$) |
| Matching | *hh* | – | *th, hh* | *th* |
| Expected nature of reaction | no reaction | no reaction | catalytic turnover | only TMSD |

| Positive Control | Concentrations in nM | |
|---|---|---|
| $S_2$-$R_1$-D | 5 | 10 |

| Reactants | Concentrations in nM | | | |
|---|---|---|---|---|
| $S_2$-L | 10 | 10 | 10 | 10 |
| $R_2$ | 10 | 10 | 10 | 10 |
| D | 10 | 10 | 10 | 10 |
| Template (added manually) | 2 ($T_{11}$) | 2 ($T_{12}$) | 2 ($T_{21}$) | 2 ($T_{22}$) |
| Matching | – | *hh* | *th* | *th, hh* |
| Expected nature of reaction | no reaction | no reaction | only TMSD | catalytic turnover |

| Positive Control | Concentrations in nM | |
|---|---|---|
| $S_2$-$R_2$-D | 5 | 10 |

**C. Specificity of the templates in a mixed pool of monomers.** We then tested the specificity of templates by adding each template individually to a mixture of monomers $S_1$, $R_1$, $S_2$ and $R_2$. The template in question was added into pre-annealed $S_1$(a)-L, $S_2$(a)-L; $R_1$(b), $R_2$ and fuel D. The reaction setup is tabulated in Table S24.

Table S24. Reaction protocol for the test of specificity for each template in a mixture of all four monomers for experiments shown in Figure 4b-c in manuscript.

| **Reactants** | **Concentrations in nM** | | | |
|---|---|---|---|---|
| $S_1$-L | 10 | 10 | 10 | 10 |
| $R_1$ | 10 | 10 | 10 | 10 |
| $S_2$-L | 10 | 10 | 10 | 10 |
| $R_2$ | 10 | 10 | 10 | 10 |
| D | 10 | 10 | 10 | 10 |
| Template (added manually) | 5 ($T_{11}$) | 5 ($T_{12}$) | 5 ($T_{21}$) | 5 ($T_{22}$) |

| **Positive Controls** | **Concentrations in nM** | |
|---|---|---|
| $S_1$-$R_1$-D | 5 | 10 |
| $S_1$-$R_2$-D | 5 | 10 |
| $S_2$-$R_1$-D | 5 | 10 |
| $S_2$-$R_2$-D | 5 | 10 |

**D. Gel electrophoresis for specificity of the templates in a mixed pool of monomers.** Individual templates were added into pre-annealed $S_1$(a)-L, $S_2$(a)-L; $R_1$(b), $R_2$ and fuel D. The reaction setup is tabulated in Table S25. Products of reactions were run in a PAGE gel alongside pre-annealed controls as described in the methods section of the manuscript.

Table S25. Reaction protocol for the PAGE test of specificity for each template in a mixture of all four monomers for experiments shown in Figures 4e in manuscript and S13. Each column shows the reagents that were added together and then loaded into the gel.

| **Reactants** | **Concentrations in nM** | | | | | | | | | |
|---|---|---|---|---|---|---|---|---|---|---|
| | Reac$^n$ | Reac$^n$ | Reac$^n$ | Reac$^n$ | Reac$^n$ | Reac$^n$ | Control | Control | Control | Control |
| $S_1$-L | 10 | 10 | 10 | 10 | 10 | 10 | – | – | – | – |
| $R_1$ | 10 | 10 | 10 | 10 | 10 | 10 | – | – | – | – |
| $S_2$-L | 10 | 10 | 10 | 10 | 10 | 10 | – | – | – | – |
| $R_2$ | 10 | 10 | 10 | 10 | 10 | 10 | – | – | – | – |
| D | – | 10 | 10 | 10 | 10 | 10 | – | – | – | – |
| $T_{11}$ | – | – | 2 | – | – | – | – | – | – | – |
| $T_{12}$ | – | – | – | 2 | – | – | – | – | – | – |
| $T_{21}$ | – | – | – | – | 2 | – | – | – | – | – |
| $T_{22}$ | – | – | – | – | – | 2 | – | – | – | – |
| $S_1$-$R_1$-D | – | – | – | – | – | – | 10 | – | – | – |
| $S_1$-$R_2$-D | – | – | – | – | – | – | – | 10 | – | – |
| $S_2$-$R_1$-D | – | – | – | – | – | – | – | – | 10 | – |
| $S_2$-$R_2$-D | – | – | – | – | – | – | – | – | – | 10 |

##### SI.2.5.6 Reactions with upstream gate circuitry

**A. Single-input YES gate.** The fuel Dn is initially bound to ancillary strand X, in solution with $S_1$(a)-L and $R_1$(a). The reaction was triggered with variable concentrations of the template $T_{11}$(a) and the input I1 was added immediately after injection, as described in Table S26.

Table S26. Reaction protocol for the catalytic templating regulated by a single-input gate system for experiments shown in Figure 6a in manuscript.

| **Reactants** | **Concentrations in nM** | | | | | | |
|---|---|---|---|---|---|---|---|
| $S_1$-L | 10 | 10 | 10 | 10 | 10 | 10 | 10 |
| $R_1$ | 10 | 10 | 10 | 10 | 10 | 10 | 10 |
| Dn-X | 10 | 10 | 10 | 10 | 10 | 10 | 10 |
| $T_{11}$ (added by injector) | 0 | 2 | 3 | 4 | 5 | 5 | 0 |
| I1 (added manually) | 10 | 10 | 10 | 10 | 10 | 0 | 0 |

| **Positive Controls** | **Concentrations in nM** | | | |
|---|---|---|---|---|
| $S_1$-$R_1$-Dn | 5 | 10 | 15 | 20 |

**B. Two-input AND gate** Solutions were prepared of $S_1$(a)-L, $R_1$(a) and and extended fuel Dn in a complex with ancillary strands E and XE. Also present is I1, except when demonstrating that the absence of I1 leads to no reaction. The reaction was triggered when either 0 nM or 10 nM of I2 and variable concentrations of template $T_{11}$(a) were injected simultaneously in the reaction mixture. The reaction protocol is tabulated in Table S27.

Table S27. Reaction protocol for the catalytic templating reaction coupled with a 2-input AND gate system for experiments shown in Figure 6b in manuscript.

| **Reactants** | **Concentrations in nM** | | | | | | | | |
|---|---|---|---|---|---|---|---|---|---|
| $S_1$-L | 10 | 10 | 10 | 10 | 10 | 10 | 10 | 10 | 10 |
| $R_1$ | 10 | 10 | 10 | 10 | 10 | 10 | 10 | 10 | 10 |
| E-Dn-XE | 10 | 10 | 10 | 10 | 10 | 10 | 10 | 10 | 10 |
| I1 | 10 | 10 | 10 | 10 | 10 | 10 | 0 | 0 | 0 |
| I2 (added by injector) | 10 | 10 | 10 | 10 | 10 | 0 | 10 | 0 | 0 |
| $T_{11}$ (added by injector) | 0 | 2 | 3 | 4 | 5 | 5 | 5 | 5 | 0 |

| **Positive Controls** | **Concentrations in nM** | | | |
|---|---|---|---|---|
| $S_1$-$R_1$-Dn | 5 | 10 | 15 | 20 |

**C. Catalytic gate** The fuel Dn is initially bound to ancillary strand X, in solution with $S_1$(a)-L and $R_1$(a) and Z. The reaction was triggered with a low concentration of $I1_M$ and variable concentrations of the template $T_{11}$. The reaction protocol is tabulated in Table S28. We also performed additional experiments with high concentrations of the sequestered fuel and monomers, as outlined in in Table S29.

Table S28. Reaction protocol for the catalytic templating coupled with a catalytic gate system for experiments shown in Figure 6c in manuscript.

| **Reactants** | | | **Concentrations in nM** | | | | |
|---|---|---|---|---|---|---|---|
| $S_1$-L | 10 | 10 | 10 | 10 | 10 | 10 | 10 |
| $R_1$ | 10 | 10 | 10 | 10 | 10 | 10 | 10 |
| Dn-X | 10 | 10 | 10 | 10 | 10 | 10 | 10 |
| Z | 10 | 10 | 10 | 10 | 10 | 10 | 10 |
| $I1_M$ (added by injector) | 2 | 2 | 2 | 2 | 2 | 0 | 0 |
| $T_{11}$ (added by injector) | 0 | 2 | 3 | 4 | 5 | 5 | 0 |

| **Positive Controls** | | **Concentrations in nM** | | |
|---|---|---|---|---|
| $S_1$-$R_1$-Dn | 5 | 10 | 15 | 20 |

Table S29. Reaction protocol for the catalytic templating coupled with a catalytic gate, with high monomers to template ratio - for experiments shown in Figure 6e in manuscript.

| **Reactants** | | | **Concentrations in nM** | | |
|---|---|---|---|---|---|
| $S_1$-L | 50 | 50 | 50 | 50 | 50 |
| $R_1$ | 50 | 50 | 50 | 50 | 50 |
| Dn-X | 100 | 100 | 100 | 100 | 100 |
| Z | 100 | 100 | 100 | 100 | 100 |
| $T_{11}$ (added by injector) | 0 | 0.5 | 1.0 | 1.5 | 2.0 |
| $I1_M$ (added manually) | 2 | 2 | 2 | 2 | 2 |

| **Positive Controls** | | **Concentrations in nM** | | |
|---|---|---|---|---|
| $S_1$-$R_1$-Dn | 5 | 10 | 50 | 100 |

**D. NOT gate** The fuel Dn is initially bound to ancillary strand X, in solution with $S_1$(a)-L and $R_1$(a) and Z. The reaction was triggered with a low concentration of $I1_M$, variable concentrations of the template $T_{11}$, and a large concentration of $I\bar{1}_M$. The precise conditions are tabulated in Table S30. Note that the catalytic experiments without $I\bar{1}_M$ function as a negative control for this experiment.

Table S30. Reaction protocol for catalytic templating coupled with a NOT gate for experiments shown in Figure 6d in manuscript.

| **Reactants** | **Concentrations in nM** | | | | |
|---|---|---|---|---|---|
| $S_1$-L | 10 | 10 | 10 | 10 | 10 |
| $R_1$ | 10 | 10 | 10 | 10 | 10 |
| Dn-X | 10 | 10 | 10 | 10 | 10 |
| Z | 10 | 10 | 10 | 10 | 10 |
| $I1_M$ (added by injector) | 2 | 2 | 2 | 2 | 2 |
| $T_{11}$ (added by injector) | 0 | 2 | 3 | 4 | 5 |
| $I\bar{1}_M$ (added manually) | 25 | 25 | 25 | 25 | 25 |

| **Positive Controls** | **Concentrations in nM** | | | |
|---|---|---|---|---|
| $S_1$-$R_1$-Dn | 5 | 10 | 15 | 20 |

### SI.3 Data processing

This section describes how raw fluorescence signals (AFU) are converted into inferred concentrations or into a normalised, dimensionless readout. Section SI.3.2 defines a plate-level normalisation based on positive controls; subsequent subsections apply assay-specific signal models and inference steps. The data processing pipeline is carried out using the in-house software Clearissa v1.71, available at `https://github.com/PrinciplesKJ/Clearissa_1.71`. Step-by-step instructions for reproducing the data processing as described in the following sections are provided in `https://doi.org/10.5281/zenodo.18674501`.

Raw fluorescence signals can exhibit time-dependent changes due to instrument drift and handling steps (for example, sealing, injections, or plate ejection and reinsertion). We assume that the measured fluorescence is the sum of contributions from the individual species present in a reaction well. Each species contribution is proportional to its concentration, and the *ratios* of the corresponding proportionality constants do not change under these perturbations.

#### SI.3.1 General experimental context and common notation

A "well" refers to an individual reaction well on a plate. Each plate contains the following:

- a buffer-only blank well with raw fluorescence $f_{\text{blank}}(t)$ (AFU),
- an assay-specific negative-control well with raw fluorescence $f_{\text{neg}}(t)$ (AFU),
- one or more positive-control wells containing a fluorescent reference species at known concentration(s) $[C]_q$, with raw fluorescence $f_{\text{pos},q}(t)$ (AFU). Here typically $[C]_q \in \{5, 10, 15, 20\}$ nM, and in some cases $\{10,\ 20,\ 50,\ 100\}$ nM for reactions with high monomer concentrations,
- experimental wells with raw fluorescence $f_{\text{raw}}(t)$ (AFU).

Here, $f_{\text{raw}}(t)$ denotes the raw fluorescence time trace of an arbitrary experimental well; when multiple experimental wells are indexed explicitly we write $f_{\text{raw},i}(t)$.

Unless stated otherwise, background correction is performed by blank subtraction:

$$f_{\text{corr}}(t) = f_{\text{raw}}(t) - f_{\text{blank}}(t), \tag{S1}$$

where $f_{\text{corr}}(t)$ is the blank-subtracted trace (AFU). The same procedure is applied to positive control and negative control traces.

###### SI.3.1.1 Definitions

- Initialisation window: a 15 min pre-trigger interval

  $$t \in [t_{\text{init,start}},\, t_{\text{init,end}}], \tag{S2}$$

  acquired while wells contain the initial 150 $\mu$L mixture (before addition of the remaining 50 $\mu$L to reach 200 $\mu$L). This window is used to estimate a mini calibration curve.

- Oversaturation window: a 15 min interval after the injection of a saturating level of a strand or complex intended to push one species to react completely

  $$t \in [t_{\text{over,start}},\, t_{\text{over,end}}]. \tag{S3}$$

  It is necessary to wait for a stable (plateau) signal before this window can be taken.

- Main measurement window: the interval used for kinetic analysis, starting at the trigger injection timepoint $t_{\text{inj}}$ and ending once the signal has reached a steady state.

  $$t \in [t_{\text{meas,start}},\, t_{\text{meas,end}}], \qquad t_{\text{meas,start}} = t_{\text{inj}}. \tag{S4}$$

Here, $t_{\text{meas,end}}$ is defined as the earliest timepoint at which the trace has reached a stable plateau; for slow reactions, $t_{\text{meas,end}}$ is set to the final recorded timepoint.

- $\text{pos}_{q,\text{init}}$: mean blank-subtracted positive-control fluorescence in the initialisation window (AFU). The index q refers to the respective known concentration of the positive control.
- $c_{\text{io}}$: fluorescence-to-concentration slope (AFU per nM) of the positive control, determined from the initialisation window.
- $C_{\text{ref}}$: reference concentration used when a concentration-referenced trajectory is required (typically 10 nM).
- $r_q(t)$: signal drift ratio for positive control $q$ (dimensionless).

#### SI.3.2 Positive control normalisation

Positive control normalisation defines a time-dependent, background-corrected reference trajectory $p_{\text{corr}}(t)$ (AFU) that corrects plate-wide, time-dependent artifacts (gain drift, handling steps, plate ejection/reinsertion).

##### SI.3.2.1 Procedure

**A. Blank subtraction.** We first correct the positive control for the background blank signal.

$$f_{\text{pos},q,\text{corr}}(t) = f_{\text{pos},q}(t) - f_{\text{blank}}(t). \tag{S5}$$

**B. Calibration slope from the initialisation window.** A mini calibration of fluorescence against concentration for the positive control is formed by linear regression of $\text{pos}_{q,\text{init}}$ against $[C]_q$:

$$\text{pos}_{q,\text{init}} = c_{\text{io}}\,[C]_q. \tag{S6}$$

We thus assume a no-intercept model after blank subtraction. For the purposes of our analysis, $c_{\mathrm{io}}$ is taken as the ground truth with which other concentrations can be converted to and from fluorescence.

**C. Plate-level drift profile.** Each positive-control trace is converted to a ratio relative to its own initialisation mean:

$$r_q(t) = \frac{\mathrm{pos}_{q,\mathrm{corr}}(t)}{\mathrm{pos}_{q,\mathrm{init}}}. \tag{S7}$$

The signal drift profile is the mean across all $Q$ positive controls:

$$r_{\mathrm{mean}}(t) = \frac{1}{Q}\sum_{q=1}^{Q} r_q(t). \tag{S8}$$

**D. Reference trajectory in AFU.** The information from all positive controls can be used to define a time-dependent nominal positive control fluorescence signal at any concentration $C_{\mathrm{ref}}$ according to:

$$p_{\mathrm{corr}}(t) = c_{\mathrm{io}}\, C_{\mathrm{ref}}\, r_{\mathrm{mean}}(t). \tag{S9}$$

Equivalently,

$$p_{\mathrm{corr}}(t) = c_{\mathrm{io}}\, C_{\mathrm{ref}}\, \frac{1}{Q}\sum_{q=1}^{Q} \frac{f_{\mathrm{pos},q}(t) - f_{\mathrm{blank}}(t)}{f_{\mathrm{pos},q,\mathrm{init}}}. \tag{S10}$$

**E. Applying the normalisation.** For an experimental well with raw fluorescence trace $f_{\mathrm{raw}}(t)$ (AFU), we first apply blank subtraction (Eq. S1) and then divide by $p_{\mathrm{corr}}(t)$ to give a ratio of the signal to the positive control:

$$g(t) = \frac{f_{\mathrm{corr}}(t)}{p_{\mathrm{corr}}(t)}. \tag{S11}$$

When concentration-referenced units are required, we define $h(t)$ with units of nM:

$$h(t) = g(t)\, C_{\mathrm{ref}}. \tag{S12}$$

$h(t)$ gives the concentration of positive control species that would be required to explain the observed fluorescence in the experimental trace.

##### SI.3.3 Conversion of AFU to concentrations for TMSD reactions (step 1)

###### SI.3.3.1 Experimental context and signal model

The assay follows step 1, a toehold-mediated strand displacement reaction of the form:

$$\mathrm{S_1\text{-}L} + \mathrm{T_{11}} \rightarrow \mathrm{S_1\text{-}T_{11}} + \mathrm{L}. \tag{S13}$$

Reaction wells were initially loaded with a fluorophore-quencher duplex $S_1$-L. The non-labelled invader strand $T_{11}$ was injected to initiate the reaction in which L is displaced and $S_1$-$T_{11}$ is formed. $S_1$-L exhibits residual fluorescence due to incomplete quenching, whereas the displacement product $S_1$-$T_{11}$ is fully fluorescent.

Positive-control wells contain $S_1$-$T_{11}$ at known concentrations, which are used to construct $p_{\mathrm{corr}}(t)$.

We apply the common preprocessing described in SI.3.2 to obtain $h(t)$ (nM) for each experimental trace (Eq. S12), i.e.

$$h(t) = \frac{f_{\mathrm{raw}}(t) - f_{\mathrm{blank}}(t)}{p_{\mathrm{corr}}(t)} C_{\mathrm{ref}}. \tag{S14}$$

For concentration inference we assume

$$h(t) = [\mathrm{S_1\text{-}T_{11}}] + \beta\,[\mathrm{S_1\text{-}L}], \tag{S15}$$

where $\beta$ is the fractional fluorescence of $S_1$-L relative to $S_1$-$T_{11}$ (dimensionless). We further assume mass conservation

$$[\mathrm{S_1\text{-}L}]_0 = [\mathrm{S_1\text{-}L}] + [\mathrm{S_1\text{-}T_{11}}], \tag{S16}$$

so that

$$h(t) = [\mathrm{S_1\text{-}T_{11}}](1-\beta) + \beta\,[\mathrm{S_1\text{-}L}]_0, \tag{S17}$$

and thus

$$[\mathrm{S_1\text{-}T_{11}}] = \frac{h(t) - \beta[\mathrm{S_1\text{-}L}]_0}{1-\beta}. \tag{S18}$$

###### SI.3.3.2 Concentration inference procedure (per trace)

**A. Determine the initial substrate concentration $[\mathrm{S_1\text{-}L}]_0$ by oversaturation.** Under oversaturation by injection of excess $\mathrm{T_{11}}$ we assume full conversion of $\mathrm{S_1}$-L to $\mathrm{S_1\text{-}T_{11}}$, hence $h(t) = [\mathrm{S_1\text{-}L}]_0$ in the oversaturated plateau window. We estimate

$$[\mathrm{S_1\text{-}L}]_0 = \mathrm{mean}(h(t)) \quad \text{for } t \in [t_{\mathrm{over,start}},\, t_{\mathrm{over,end}}]. \tag{S19}$$

**B. Estimate $\beta$ from the pre-trigger window.** In the initialisation window, $[\mathrm{S_1\text{-}T_{11}}] = 0$ and $h(t) = \beta[\mathrm{S_1\text{-}L}]_0$. We compute

$$h_{\mathrm{init}} = \mathrm{mean}(h(t)) \quad \text{for } t \in [t_{\mathrm{init,start}},\, t_{\mathrm{init,end}}], \tag{S20}$$

and estimate

$$\beta = \frac{h_{\mathrm{init}}}{[\mathrm{S_1\text{-}L}]_0}. \tag{S21}$$

This estimate of $\beta$ is calculated separately for each reaction well.

**C. Reconstruct concentrations over time** Having identified the constants $[S_1\text{-}L]_0$ and $\beta$, Eq. S18 can be used to infer the concentrations of the product $[\mathrm{S_1\text{-}T_{11}}]$ over time.

#### SI.3.4 Conversion of AFU to concentrations for HMSD reactions (step 2)

##### SI.3.4.1 Experimental context and signal model

HMSD (step 2) was monitored in the donor channel of a donor-acceptor FRET pair. The donor-labelled complex $S_1$-$T_{11}$ was initially present in solution and provides the bright donor reference signal. Upon a HMSD reaction with $R_1$, a ternary complex forms in which donor emission was quenched by FRET:

$$S_1\text{-}T_{11} + R_1 \rightarrow S_1\text{-}T_{11}\text{-}R_1. \tag{S22}$$

Consequently, the donor-channel fluorescence decreases as product accumulates.

We initially considered inferring product formation from the FRET readout; however, in practice the donor-channel signal provided a stronger signal-to-noise ratio and is therefore used for concentration inference.

A donor reference control containing pre-annealed $S_1$-$T_{11}$ at a known concentration of 10 nM was used for donor-channel normalisation. Note that this control contains 0 nM product and is therefore not the positive control in the sense it is in the TMSD conversion model. The donor reference control serves as the normalisation reference rather than as a baseline to be subtracted, and is therefore not plotted alongside the experimental product traces.

Because only a single $S_1$-$T_{11}$ reference trace is available, $h(t)$ is obtained by blank subtraction followed by pointwise normalisation to the blank-subtracted $S_1$-$T_{11}$ reference trajectory and scaling by its known concentration.

We apply blank subtraction in the donor channel to obtain

$$f_{\text{corr}}(t) = f_{\text{raw}}(t) - f_{\text{blank}}(t), \qquad f_{\text{ref,corr}}(t) = f_{\text{ref}}(t) - f_{\text{blank}}(t), \tag{S23}$$

where $f_{\mathrm{raw}}(t)$ is the experimental donor-channel trace and $f_{\mathrm{ref}}(t)$ is the donor reference-control trace (pre-annealed $S_1$-$T_{11}$). In the HMSD workflow, $f_{\mathrm{ref,corr}}(t)$ serves the same functional role as $p_{\mathrm{corr}}(t)$ in the TMSD conversion, but is taken directly from a single donor reference control rather than being constructed from multiple positive-control wells. We then form the pointwise ratio

$$g(t) = \frac{f_{\mathrm{corr}}(t)}{f_{\mathrm{ref,corr}}(t)}, \tag{S24}$$

and define the concentration-referenced donor signal

$$h(t) = g(t)\,[\mathrm{S_1\text{-}T_{11}}]_{\mathrm{ref}}, \tag{S25}$$

where $[\mathrm{S_1\text{-}T_{11}}]_{\mathrm{ref}}$ is the known concentration of the donor reference control.

Concentration inference uses the linear signal model

$$h(t) = [\mathrm{S_1\text{-}T_{11}}](t) + \beta\,[\mathrm{S_1\text{-}T_{11}\text{-}R_1}](t), \tag{S26}$$

where $\beta$ is the residual donor brightness of the product $S_1$-$T_{11}$-$R_1$ relative to $S_1$-$T_{11}$. Assuming conservation of the donor-labelled complex,

$$[\mathrm{S_1\text{-}T_{11}}]_0 = [\mathrm{S_1\text{-}T_{11}}](t) + [\mathrm{S_1\text{-}T_{11}\text{-}R_1}](t), \tag{S27}$$

we obtain

$$[\mathrm{S_1\text{-}T_{11}\text{-}R_1}](t) = \frac{[\mathrm{S_1\text{-}T_{11}}]_0 - h(t)}{1-\beta}. \tag{S28}$$

###### SI.3.4.2 Concentration inference procedure

Compared to the TMSD concentration inference procedure (Sec. SI.3.3), HMSD differs in three respects: (i) product formation decreases the donor-channel fluorescence due to quenching; (ii) the donor reference control is a single-concentration $S_1$-$T_{11}$ trace used for pointwise normalisation; and (iii) the quenching factor $\beta$ is estimated from separate pre-annealed $S_1$-$T_{11}$ and $S_1$-$T_{11}$-$R_1$ control wells.

**A. Estimate $\beta$ from control wells.** The residual brightness factor $\beta$ is estimated from donor-channel measurements of pre-annealed control complexes. The unquenched donor reference control $S_1$-$T_{11}$ is typically measured in a single well at a known concentration (typically $[S_1\text{-}T_{11}]_{\text{ref}} = 10$ nM); after blank subtraction its donor-channel signal is denoted $f_{\text{ref,corr}}(t)$. The quenched product control $S_1$-$T_{11}$-$R_1$ is typically measured in four wells at different concentrations and is processed using the positive control normalisation procedure (Sec. SI.3.2) to yield a drift-corrected reference trajectory in the donor channel, denoted $p_{\text{corr,prod}}(t)$.

We compute $\beta$ from the ratio of initialisation-window means,

$$\beta = \frac{\text{mean}(p_{\text{corr,prod}}(t))}{\text{mean}(f_{\text{ref,corr}}(t))} \quad \text{for } t \in [t_{\text{init,start}},\, t_{\text{init,end}}]. \tag{S29}$$

Because both $p_{\text{corr,prod}}(t)$ and $f_{\text{ref,corr}}(t)$ are expressed on the same concentration-referenced scale (using the same reference concentration, typically 10 nM), the choice of reference concentration cancels in the ratio and $\beta$ is treated as concentration-independent. For a given plate (experiment), the resulting $\beta$ is treated as a fixed constant and applied to all experimental wells on that plate.

**B. Estimate $[S_1\text{-}T_{11}]_0$ and reconstruct product concentrations.** Prior to reaction initiation, no product has formed, so $[S_1\text{-}T_{11}\text{-}R_1](t) = 0$ and therefore $h(t) = [S_1\text{-}T_{11}]_0$ in the initialisation window. We estimate

$$[S_1\text{-}T_{11}]_0 = \text{mean}(h(t)) \quad \text{for } t \in [t_{\text{init,start}},\, t_{\text{init,end}}], \tag{S30}$$

and then use Eq. S28 to infer $[\mathrm{S_1\text{-}T_{11}\text{-}R_1}](t)$. Note that $[\mathrm{S_1\text{-}T_{11}}]_0$ is calculated individually for every trace. Due to measurement noise, inferred product concentrations slightly below zero (down to $-1$ nM) are permitted.

##### SI.3.5 Conversion of AFU to concentrations for internal TMSD reactions (step 3)

###### SI.3.5.1 Experimental context and signal model

Internal TMSD follows:

$$\mathrm{D} + \mathrm{S_1\text{-}T_{11}\text{-}R_1} \rightarrow \mathrm{S_1\text{-}R_1\text{-}D} + \mathrm{T_{11}}. \tag{S31}$$

Fuel D was pre-incubated in the wells before the reaction starts, and $\mathrm{S_1\text{-}T_{11}\text{-}R_1}$ is injected at time $t_{\mathrm{inj}}$. The injection produces an instantaneous fluorescence jump due to residual $\mathrm{S_1\text{-}T_{11}\text{-}R_1}$ fluorescence (incomplete quenching). This injection offset is corrected using a negative control well containing $\mathrm{S_1\text{-}T_{11}\text{-}R_1}$ only (no fuel).

Positive-control wells contain the fully fluorescent product species $\mathrm{S_1\text{-}R_1\text{-}D}$ at known concentrations, and these traces are used to construct $p_{\mathrm{corr}}(t)$ for the channel analysed here (Sec. SI.3.2).

We apply the same procedure as in SI.3.3, namely assuming

$$[\mathrm{S_1\text{-}R_1\text{-}D}] = \frac{h(t) - \beta[\mathrm{S_1\text{-}T_{11}\text{-}R_1}]_0}{1-\beta}, \tag{S32}$$

and estimating the constants $\beta$ and $[\mathrm{S_1\text{-}T_{11}\text{-}R_1}]_0$. $[\mathrm{S_1\text{-}T_{11}\text{-}R_1}]_0$ can be inferred from the signal after oversaturation with excess D, which drives complete conversion of $\mathrm{S_1\text{-}T_{11}\text{-}R_1}$ to $\mathrm{S_1\text{-}R_1\text{-}D}$ in each experimental well.

$$[\mathrm{S_1\text{-}T_{11}\text{-}R_1}]_0 = \mathrm{mean}(h_{\mathrm{exp}}(t)) \quad \text{for } t \in [t_{\mathrm{over,start}},\, t_{\mathrm{over,end}}]. \tag{S33}$$

However, the initial time points cannot be used to estimate $\beta$ because the $\mathrm{S_1\text{-}T_{11}\text{-}R_1}$ complex was not in solution prior to injection. We therefore use a negative control well (containing $\mathrm{S_1\text{-}T_{11}\text{-}R_1}$ but no D), in which no product forms and thus $h_{\mathrm{neg}}(t) = \beta[\mathrm{S_1\text{-}T_{11}\text{-}R_1}]_0$. We define

$$h_{\mathrm{neg,base}} = \mathrm{mean}(h_{\mathrm{neg}}(t)) \quad \text{for } t \in [t_{\mathrm{nc,start}},\, t_{\mathrm{nc,end}}], \tag{S34}$$

where $[t_{\mathrm{nc,start}},\, t_{\mathrm{nc,end}}]$ refers to a 30 min interval selected after signal stabilisation in the negative-control trace.

To obtain a single $\beta$ for the experiment, we use a robust plate-level estimate of the initial substrate concentration, $[\mathrm{S_1\text{-}T_{11}\text{-}R_1}]_{0,\mathrm{plate}}$, defined as the median of $[\mathrm{S_1\text{-}T_{11}\text{-}R_1}]_{0,\mathrm{well}}$ across experimental wells. A single value of $\beta$ is then estimated as

$$\beta = \frac{h_{\mathrm{neg,base}}}{[\mathrm{S_1\text{-}T_{11}\text{-}R_1}]_{0,\mathrm{plate}}}, \tag{S35}$$

and this value is applied to all experimental wells for the experiment. This differs from the standard TMSD analysis, where $\beta$ is estimated separately for each reaction well.

Having identified the constants $[S_1\text{-}T_{11}\text{-}R_1]_0$ per well and a global $\beta$, Eq. S32 can be used to infer the concentrations of the product $[S_1\text{-}R_1\text{-}D]$ over time.

#### SI.3.6 Calculation of a control-normalised readout in relative units

##### SI.3.6.1 Experimental context

This mode reports a dimensionless normalised readout (relative fluorescence, RF) rather than concentrations. It is used when a fluorescence-to-concentration mapping is not justified (for example, complex stoichiometry, multiple fluorescent species, or unknown quenching fractions). The essential principle is to report the fluorescence on a scale normalised with respect to two controls – typically a negative control assigned the value 0 and a positive control assigned the value 1. Note that the system being reported need not contain only the fluorescent species that define these controls.

For each well, we consider the blank-corrected fluorescence $f_{\text{corr}}(t)$ (Eq. S1) and the averaged, blank-corrected positive control $p_{\text{corr}}(t)$ (Eq. S9). We also consider a blank-corrected negative control

$$f_{\text{neg,corr}}(t) = f_{\text{neg}}(t) - f_{\text{blank}}(t). \tag{S36}$$

All quantities are defined for the time period corresponding to the main measurement interval

$$t \in [t_{\text{meas,start}},\ t_{\text{meas,end}}]. \tag{S37}$$

**A. Negative-control baseline (AFU).** A (typically 30 min) baseline window $t \in [t_{\text{nc,start}}, , t_{\text{nc,end}}]$ within the main measurement interval $t \in [t_{\text{meas,start}}, , t_{\text{meas,end}}]$ is selected after any initial transients in the negative control due to the reaction initiation procedure have subsided. We use this time period to define a fixed negative control baseline

$$f_{\text{neg,base}} = \text{mean}(f_{\text{neg,corr}}(t)) \quad \text{for } t \in [t_{\text{nc,start}},\ t_{\text{nc,end}}]. \tag{S38}$$

**B. Relative fluorescence calculation.** We then calculate a relative fluorescence as

$$\text{RF}(t) = \frac{f_{\text{corr}}(t) - f_{\text{neg,base}}}{p_{\text{corr}}(t) - f_{\text{neg,base}}}. \tag{S39}$$

Note that this approach ignores (typically very small) changes in the negative control signal over the main measurement window $t \in [t_{\text{meas,start}},\ t_{\text{meas,end}}]$.

### SI.4 Kinetic model fitting

This section describes how the inferred concentration traces from our experiments are fitted to kinetic models. The data fitting pipeline is performed using in-house software Clearissa v1.71, available at `https://github.com/PrinciplesKJ/Clearissa_1.71`. The specific step by step instructions to reproduce the fits are outlined in `https://doi.org/10.5281/zenodo.18674501`.

#### SI.4.1 Single step bimolecular reactions

This section describes how concentration trajectories for TMSD, internal toehold TMSD and HMSD,inferred from fluorescence (Secs. SI.3.3, SI.3.5 and SI.3.4) are fitted to a standard mass-action model. Fits are performed only over a user-defined kinetic window $t \in [t_0, t_{\mathrm{meas,end}}]$, where $t_0$ is the reaction-onset time chosen in the analysis software, typically aligned to the trigger injection and excluding any pre-onset baseline. The model assumes well-mixed mass-action kinetics and constant volume over the fitted window.

All single-step reactions are modelled as an irreversible bimolecular conversion of the form

$$\mathrm{X} + \mathrm{Z} \rightarrow \mathrm{Y} + \mathrm{W}, \tag{S40}$$

where Y is the product species concentration trajectory to be fitted. W is a placeholder for any byproduct(s) that are created within the reaction but are not included in the fitted rate law. The fitted observable is $y(t) = [\mathrm{Y}](t)$.

The ordinary differential equation governing all three reactions is expected to be of the form

$$\frac{d}{dt}[\mathrm{Y}](t) = +k_f[\mathrm{X}](t)[\mathrm{Z}](t), \tag{S41}$$

where Y is the product and X and Z are the reactants. The conservation of mass

$$[\mathrm{X}](t) = [\mathrm{X}]_0 - [\mathrm{Y}](t), \tag{S42}$$

$$[\mathrm{Z}](t) = [\mathrm{Z}]_0 - [\mathrm{Y}](t), \tag{S43}$$

also applies. Substituting these conservation laws yields the reduced one-dimensional form used for fitting,

$$\frac{d}{dt}[\mathrm{Y}](t) = k_f\big([\mathrm{X}]_0 - [\mathrm{Y}](t)\big)\big([\mathrm{Z}]_0 - [\mathrm{Y}](t)\big). \tag{S44}$$

For each experimental trace, we fit a modelled trajectory $y_{\mathrm{sim}}(t)$ to the experimentally inferred $[\mathrm{Y}](t)$, using $k_f$ and $[\mathrm{X}]_0$ as fitting parameters and using the value of $[\mathrm{Z}]_0$ calculated when inferring $[\mathrm{Y}](t)$.

###### SI.4.1.1 Mapping to assay-specific species.

Below, we state which species corresponds to $X, Y, Z$ in each experiment that is fitted.

##### A. Standard TMSD (Sec. SI.3.3).

$$\mathrm{X} \equiv \mathrm{T}_{11}, \qquad \mathrm{Z} \equiv \mathrm{S}_1\text{-L}, \qquad \mathrm{Y} \equiv \mathrm{S}_1\text{-T}_{11}, \qquad \mathrm{W} \equiv \mathrm{L}, \qquad y(t) = [\mathrm{S}_1\text{-T}_{11}](t). \tag{S45}$$

##### B. HMSD (Sec. SI.3.4).

$$\mathrm{X} \equiv \mathrm{R}_1, \qquad \mathrm{Z} \equiv \mathrm{S}_1\text{-T}_{11}, \qquad \mathrm{Y} \equiv \mathrm{S}_1\text{-T}_{11}\text{-R}_1, \qquad \mathrm{W} \equiv \emptyset, \qquad y(t) = [\mathrm{S}_1\text{-T}_{11}\text{-R}_1](t). \tag{S46}$$

##### C. Internal TMSD (Sec. SI.3.5).

$$\mathrm{X} \equiv \mathrm{D}, \quad \mathrm{Z} \equiv \mathrm{S_1\text{-}T_{11}\text{-}R_1}, \quad \mathrm{Y} \equiv \mathrm{S_1\text{-}R_1\text{-}D}, \quad \mathrm{W} \equiv \mathrm{T_{11}}, \quad y(t) = [\mathrm{S_1\text{-}R_1\text{-}D}](t). \tag{S47}$$

###### SI.4.1.2 Fitting procedure

Fits are performed over the fitting window $t \in [t_0, t_{\text{meas,end}}]$, where $t_0 > t_{\text{meas,start}} = t_{\text{inj}}$ is the reaction-onset time chosen in the analysis software (typically aligned to the trigger injection and excluding any pre-onset baseline). For a candidate parameter set $k_f$, $[X]_0$ and fixed $[Z]_0$, the model prediction is evaluated using the analytical solution of the reduced mass-action model. Parameter estimation is performed by nonlinear least squares in Python using `scipy.optimize.curve_fit` with non-negativity bounds, fitting $y_{\text{sim}}(t)$ to the inferred product trace $[\mathrm{Y}](t)$ over the fitted time points. The default initial guess for the rate constant is $k_f = 10^5\ \mathrm{M^{-1}s^{-1}}$, and the initial guess for $[\mathrm{X}]_0$ is 10 nM. Fit quality is quantified using the coefficient of determination $R^2$, computed from the residual sum of squares between $[\mathrm{Y}](t)$ and $y_{\text{sim}}(t)$, normalised by the total sum of squares of $[\mathrm{Y}](t)$ about its mean over the fitted time points.

###### SI.4.1.3 Infererence of a representative rate constant from replicate fits.

Replicate traces sharing the same nominal initial conditions are grouped together. Each trace in a group is fitted individually, yielding a rate constant $k_{f,i}$ and fitted concentration $X_{0,i}$ for trace $i$. The fixed concentration $[Z]_{0,i}$ (obtained during concentration inference) may also vary slightly across replicates due to experimental variations.

For each group, the representative rate constant is taken as the arithmetic mean of the replicate values $k_{f,i}$. Representative concentrations are defined analogously as the arithmetic means of $[X]_{0,i}$ and $[Z]_{0,i}$ across replicates.

For visualisation, we plot the group mean experimental trajectory and a shaded envelope spanning the pointwise minimum and maximum across replicate traces. The group fit trajectory is obtained by simulating the kinetic model using the averaged parameters.

#### SI.4.2 Kinetic model fitting of catalytic turnover experiments

We fit the normalized fluorescence in Figures 3a-d in the manuscript using a very rough model that assumes: (a) that the normalized fluorescence is proportional the concentration of product (we do not distinguish between template-bound and free in solution here); and (b) the template reaches a quasi-steady state in which it operates in a Michaelis-Menten-like way, with a turnover per template given by

$$\frac{d}{dt}[\mathrm{S_1\text{-}R_1\text{-}D}](t) = \frac{k\,[\mathrm{S_1\text{-}L}]^n}{K + [\mathrm{S_1\text{-}L}]^n}\,[T], \tag{S48}$$

where $n$, $k$ and $K$ are constants and $[T]$ is the template concentration. Here, $[\mathrm{S_1\text{-}L}]$ is being used as a stand-in for any/all the components of the product, since $[\mathrm{S_1\text{-}L}] \approx [\mathrm{S}] \approx [\mathrm{D}]$ in this particular setup.

We expect that the reaction proceeds in a step-by-step fashion, with first $S_1$ binding to $T_{11}$, then to $R_1$, and finally to D forming the completed structure. Each of these steps should be linear in the concentration of the reactants, and therefore we assume that $n = 1$ henceforth, even though the product has three components.

When the templates were injected into the solution, there is a burst of $S_1$ binding to $T_{11}$ and becoming fluorescent, before the system settles down to a state in which subsequent fluorescence increases require full catalytic turnover. This initial burst is not captured by the model in Eq. S48. Therefore, fits are performed only after a manually chosen start timepoint $t_0$ at which the trace transitions from the burst regime into a slower regime in which growth is driven by turnover, consistent with the model in Eq. S48. Within a given experiment, the same $t_0$ is applied to all traces (common reaction onset), while $t_0$ may differ between experiments.

###### SI.4.2.1 Fitting procedure

For each trace, we take the relative fluorescence RF($t$) and convert it to an estimated concentration [$S_1$-$R_1$-D]($t$), by multiplying by the (assumed known) reference concentration

$$[\mathrm{S_1\text{-}R_1\text{-}D}](t) = \mathrm{RF}(t) C_{\mathrm{ref}}. \tag{S49}$$

Template concentrations $[T]$ are assumed to match the intended value $(0, 2, 3, 4, 5$ nM) for each trace. The fitting window is $[t_0, t_{\mathrm{meas,end}}]$, where $t_0$ is chosen by inspection. The initial condition is set from the measured value at the first time point within the fitting window

$$y_{\mathrm{sim}}(t_0) = [\mathrm{S_1\text{-}R_1\text{-}D}](t_0), \tag{S50}$$

and is not fitted.

For each system (internal toehold length), we simultaneously fit all traces (including replicates) at all concentrations of template. We use a single $k$ and $K$ parameter shared for all traces, but each trace has its own fitted initial pool of building blocks $[\mathrm{S_1\text{-}L}]_0$. We assume mass conservation for the building-block pool, so that the available pool at time $t$ is $[\mathrm{S_1\text{-}L}]_0 - y_{\mathrm{sim}}(t)$.

For a candidate parameter set, the ODE

$$\frac{d}{dt} y_{\mathrm{sim}}(t) = \frac{k \left([\mathrm{S_1\text{-}L}]_0 - y_{\mathrm{sim}}(t)\right)}{K + \left([\mathrm{S_1\text{-}L}]_0 - y_{\mathrm{sim}}(t)\right)} [T] \tag{S51}$$

with initial condition $y_{\mathrm{sim}}(t_0) = [\mathrm{S_1\text{-}R_1\text{-}D}](t_0)$ is numerically integrated for each trace over $[t_0, t_{\mathrm{meas,end}}]$. Numerical integration is performed in Python using *scipy.integrate.odeint*

(LSODA).

Parameter estimation is performed by nonlinear least squares in Python using `scipy.optimize.least_squares` by minimising the residuals between $y_{\mathrm{sim},i}(t)$ and $[\mathrm{S_1\text{-}R_1\text{-}D}](t)$ over the fitted time points. Residuals are computed per trace on its fitted time grid and concatenated across traces for simultaneous optimisation. All fitted parameters are constrained to be non-negative.

Initial guesses are $k = 1\ \mathrm{min}^{-1}$, $K = 10$ nM, and $[\mathrm{S_1\text{-}L}]_0 = 10$ nM for each trace. The boundaries set for fitting are $k \in [10^{-10}, 1]\ \mathrm{min}^{-1}$, $K \in [0.5, 500]$ nM, and $[\mathrm{S_1\text{-}L}]_0$ within $\pm 20\%$ of the initial guess. These bounds were chosen to span plausible ranges given the experimental conditions.

For each system, we simultaneously fit all traces (including replicates) at all template concentrations. All traces within a fit share a common fitting window $[t_0, t_{\mathrm{meas,end}}]$, a single $k$ and $K$, while each trace has its own fitted $[\mathrm{S_1\text{-}L}]_0$.

Fit quality is quantified using the standard sum of squared residuals (least-squares error) over the fitted time points, and the corresponding fitted trajectories are inspected against the experimental traces.

The kinetic fits for the variants of $D^{\mathrm{j}}, j = 5, 6, 10$ indicate that only the ratio $k/K$ is well constrained, consistent with a reaction that operates in a non-saturating regime in which $s(t) = ([\mathrm{S_1\text{-}L}]_0 - y_{\mathrm{sim}}(t)) \ll K$ over the fitted window. For these systems. we therefore reduced Eq. S51 to a linear form to obtain

$$\frac{d}{dt} y_{\mathrm{sim}}(t) = k_{\mathrm{eff}} \left([\mathrm{S_1\text{-}L}]_0 - y_{\mathrm{sim}}(t)\right)[T], \qquad k_{\mathrm{eff}} \equiv \frac{k}{K}, \tag{S52}$$

which removes the $k$–$K$ degeneracy. For these variants only, we therefore report fits in terms of the effective rate constant $k_{\mathrm{eff}}$.

#### SI.5 Supplementary results

##### SI.5.1 Individual steps

###### SI.5.1.1 Step 1 – toehold-mediated strand displacement

Data for TMSD using a 6 nt toehold is shown in Figure S2a, alongside data corresponding to a 7 nt toehold in Figure S2b. Both datasets are obtained from single replicas.

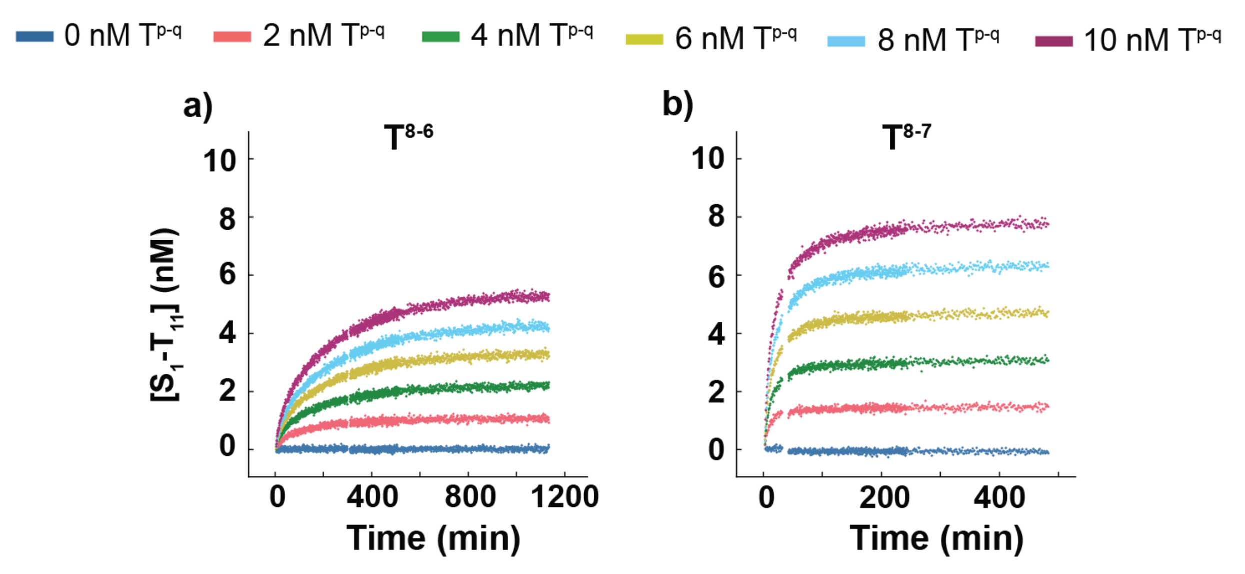

Figure S2. Reaction progress against time for TMSD with varying lengths of toehold. Reactions were conducted according to the experimental protocol outlined in Table S12, with $T^{p\text{-}q}$ added to a solution of $S_1$-L. Fluorescence in the ATTO 488 channel is converted into product concentration as described in section SI.3.3.

###### SI.5.1.2 Step 2 – handhold-mediated strand displacement

Data for HMSD using a 0 and 7-9 nt handhold is shown in Figure S3. All datasets are obtained from single replicas.

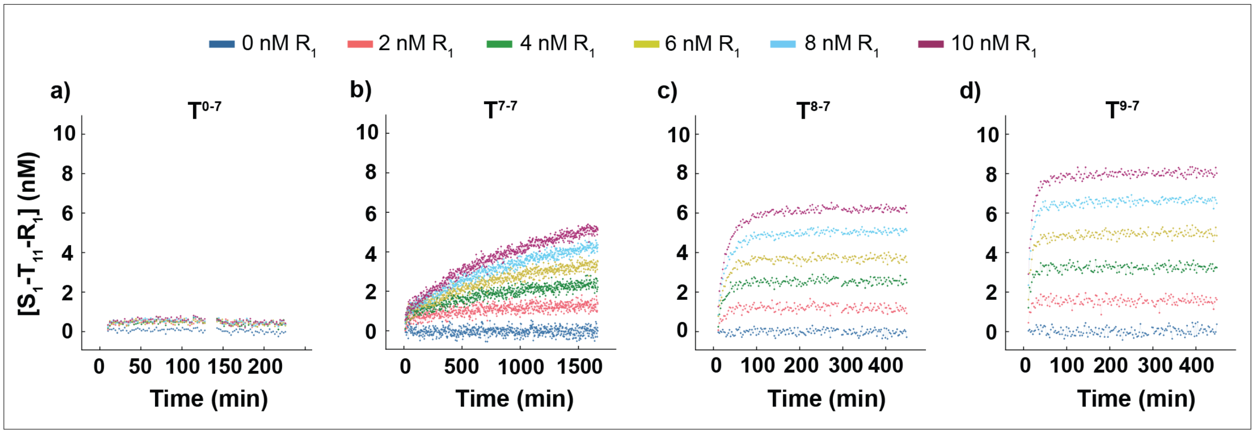

Figure S3. Reaction progress against time for HMSD with varying lengths of handhold. Reactions were conducted according to the experimental protocol outlined in Table S13, with $R_1$ added to a solution of $S_1$-$T^{p\text{-}q}$. Fluorescence in the donor channel ATTO 488 is converted into product concentration as described in section SI.3.4.

###### SI.5.1.3 Step 3 – internal toehold-mediated strand displacement

In addition to the data in Figure 2c in the manuscript, data for internal TMSD using 5, 6 and 10 nt internal toehold are shown in Figure S4a, b and c respectively. All datasets are obtained from three replicas with the mean and range plotted in each case.

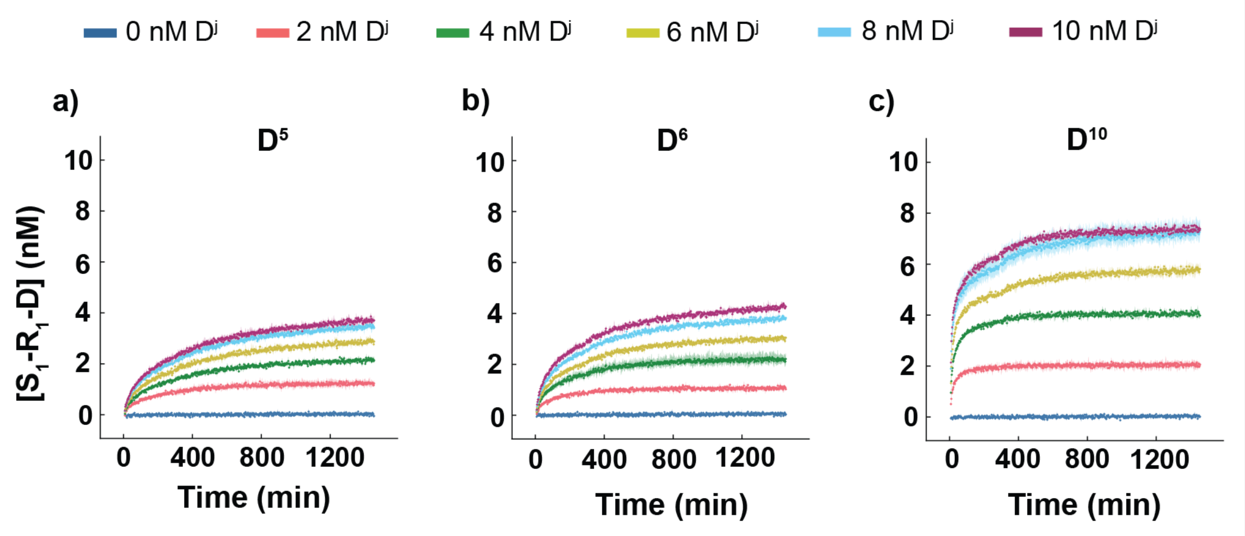

Figure S4. Reaction progress against time for internal TMSD with varying lengths of internal toehold. Reactions were conducted according to the experimental protocol outlined in Table S14, with $S_1$-$R_1$-$T_{11}$ added to a solution of $D^j$. Fluorescence in ATTO 488 channel is converted into product concentration as described in section SI.3.5.

##### SI.5.2 Test for reversibility

###### SI.5.2.1 Reversibility of step 1

Data from a single replica, probing the reversibility of step 1 is shown in Figure S5. From the graph we observe negligible differences relative to the 0 nM control, suggesting that the reaction is functionally irreversible. The jumps in the fluorescence signal were due to the change of mechanical environment during injection and applying the seal on the plate.

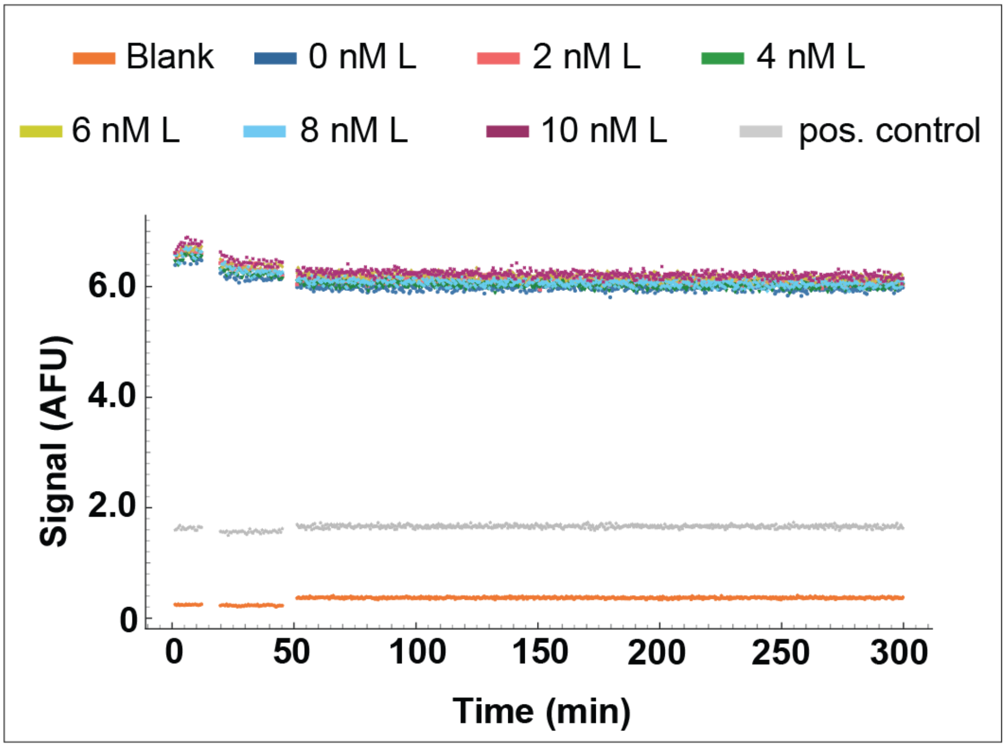

Figure S5. Reversibility test for step 1. Reactions were conducted according to the experimental protocol outlined in Table S15, with L added to a solution of $S_1$-$T_{11}$. The fluorescence in the ATTO 488 channel is plotted against time, with the 10 nM of $S_1$-L positive control experiment reported.

##### SI.5.2.2 Reversibility of step 3

Data from single replicas, probing the reversibility of step 3 is shown in Figure S6 and S7. From the graphs we observe negligible differences relative to the 0 nM control, suggesting that the reaction is functionally irreversible. The jumps in the fluorescence signal were due to the change of mechanical environment during injection and applying the seal on the plate.

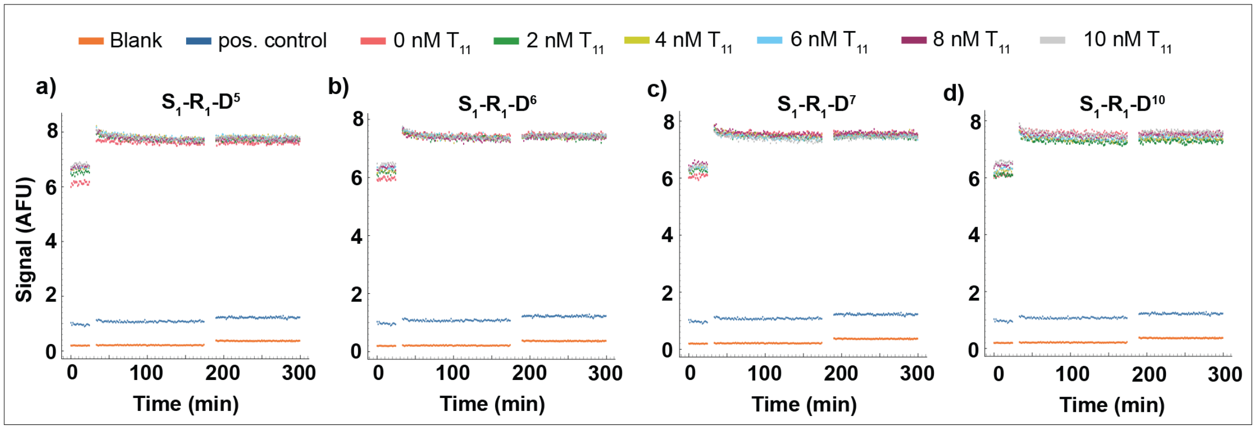

Figure S6. Reversibility test for step 3 for varying internal toeholds. Reactions were conducted according to the experimental protocol outlined in Table S16, with template $T_{11}$ added to $S_1$-$R_1$-$D^j$, (j = 5, 6, 7, 10). The fluorescence in the ATTO 488 channel is plotted against time, with the 10 nM of $S_1$-$R_1$-$T_{11}$ positive control experiment reported for comparison. No reaction was observed.

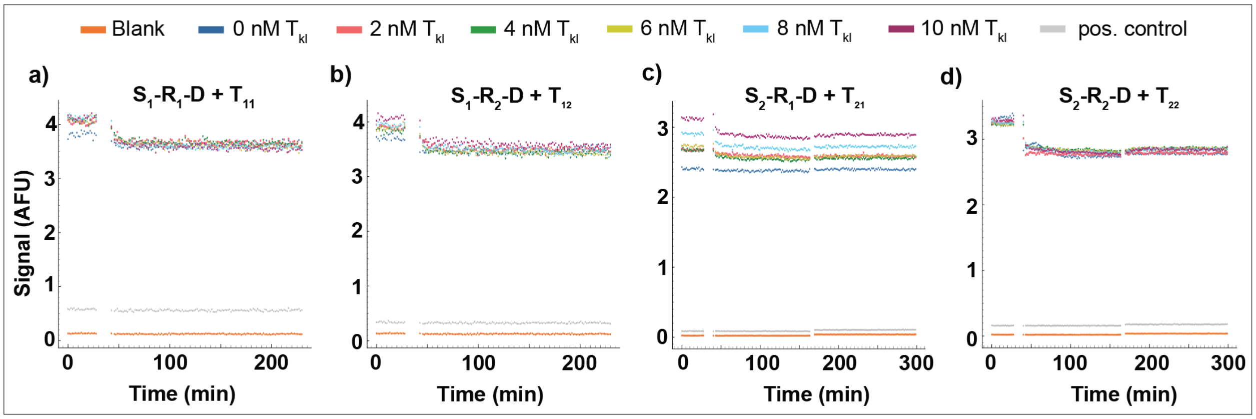

Figure S7. Reversibility test for step 3 for all orthogonal products. Reactions were conducted according to the experimental protocol outlined in Table S16, with template $T_{kl}$ added to $S_k$-$R_l$-$D^7$. The fluorescence in the ATTO 488 channel (a-b) and Alexa Fluor 546 channel (c-d) are plotted against time, with the 10 nM of $S_k$-$R_l$-$T_{kl}$ positive control experiment reported for comparison. No reaction was observed.

##### SI.5.3 Optimization of catalytic templating turnover

###### SI.5.3.1 Optimization of the toehold and the handhold lengths

Data from single replicas, probing catalytic turnover for different toehold and handhold lengths for an internal toehold of length 7 is plotted in Figure S8. We observe that while most systems show evidence of catalytic turnover, the template with toehold 7 and handhold 8 was the most efficient. From this comparative data, we observed that making the recognition domains longer could potentially slow down the overall turnover, presumably due to product inhibition.

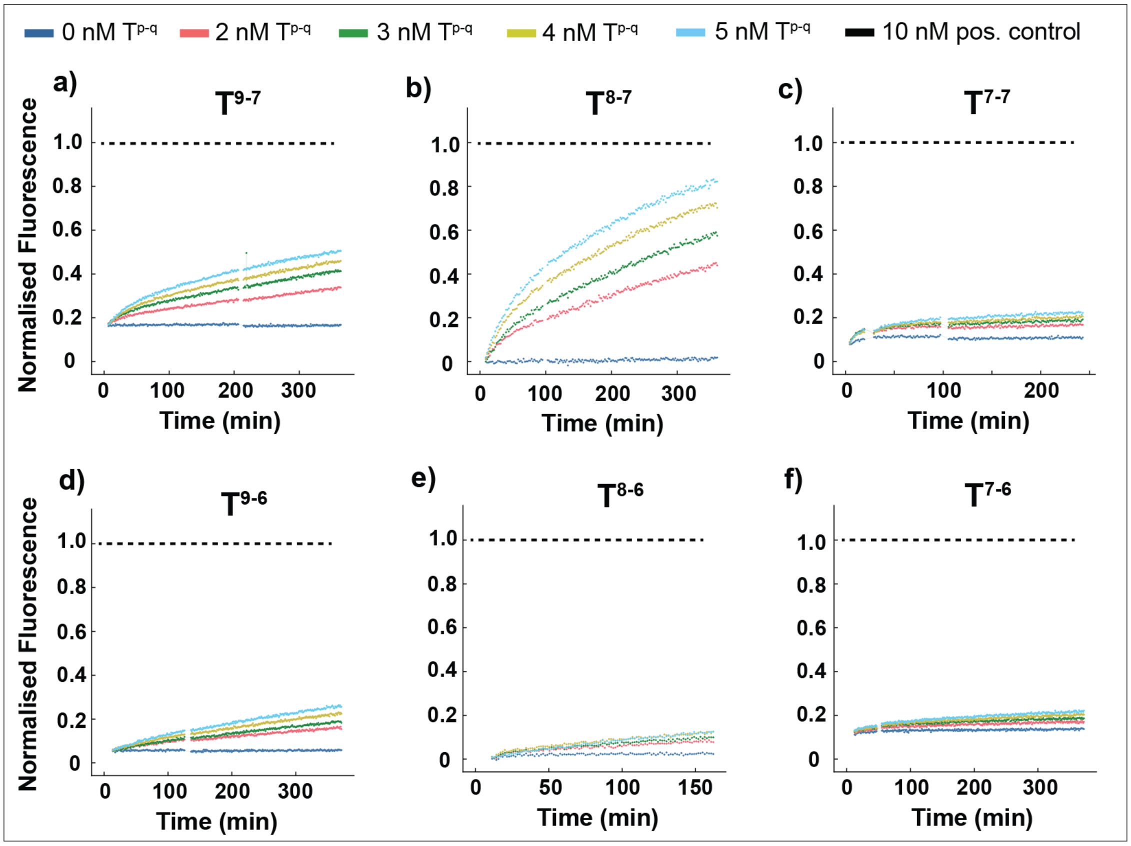

Figure S8. Catalytic turnover with variable lengths of toeholds and handholds. Reactions were conducted according to the experimental protocol outlined in Table S17, with template $T^{p\text{-}q}$ added to a solution of $S_1$-L, $R_1$ and D. The FRET in the ATTO 488-Alexa Fluor 546 (FRET) channel is plotted against time, with the 10 nM of $S_1$-$R_1$-$D^7$ positive control experiment reported for comparison. The normalised data are obtained by following the procedure in section SI.3.2.

##### SI.5.4 Catalytic templating regulated by excess of fuel molecule

In addition to the data in Figure 3e in the manuscript, we collected further data from three replicas for varying concentrations of $D^7$ and $D^6$, with the mean and range plotted in Figure S9.

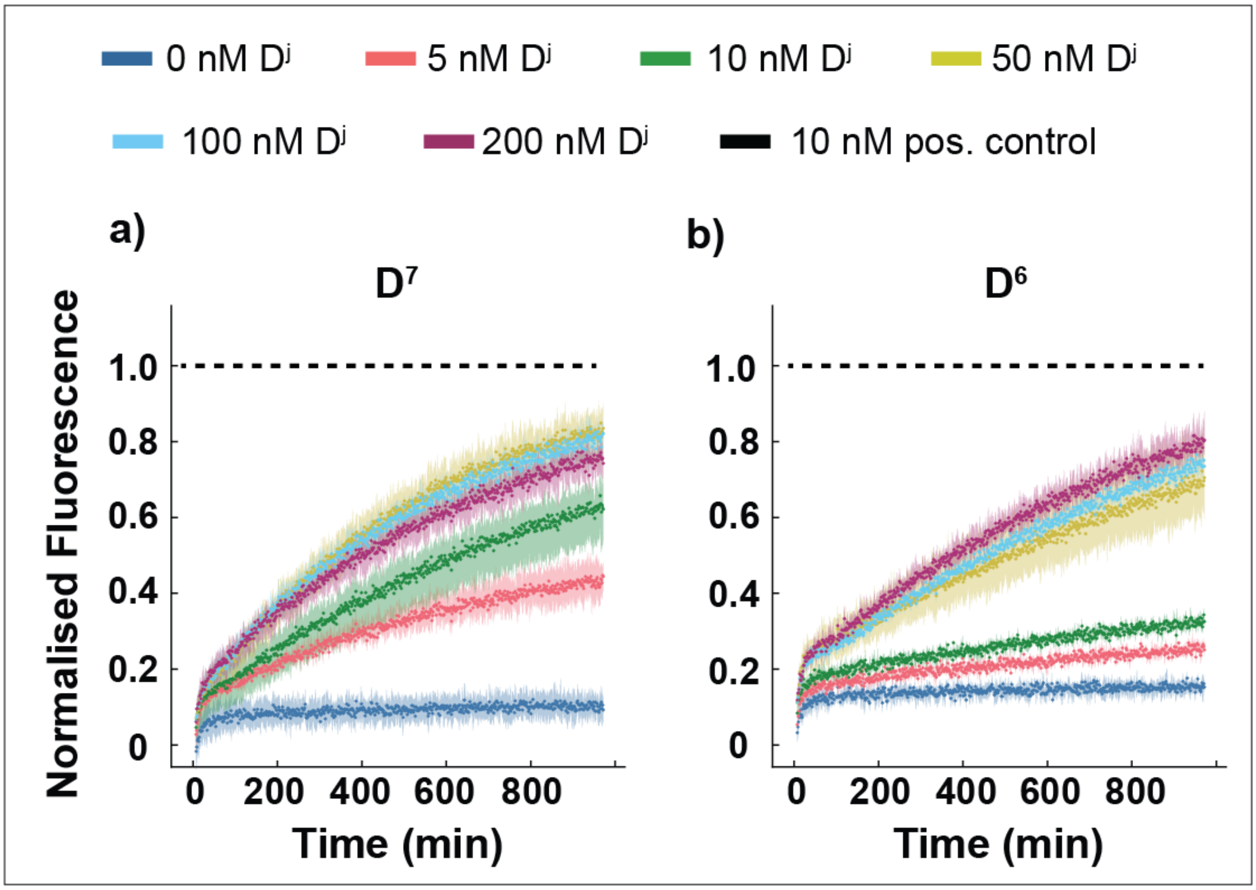

Figure S9. Catalytic turnover with varying concentration of fuels: (a) $D^7$; and (b) $D^6$. Reactions were conducted according to the experimental protocol outlined in Table S19, with template $T_{11}$ added to $S_l$-L, $R_1$ and $D^7/D^6$. The fluorescence in the ATTO 488 channel, normalized with respect 10 nM of $S_1$-$R_1$-$D^{6/7}$ according to the procedure in section SI.3.2, is plotted against time.

##### SI.5.5 Catalytic templating with high monomer-to-template ratio

In addition to the data in Figure 3f in the manuscript, we collected a third replica which showed anomalous behaviour (most likely due to formation of a bubble) in one of the traces (shown in light blue). For completeness, we report that replica in Figure S10a. In Figure S10b we show the combined data for all three replicas.

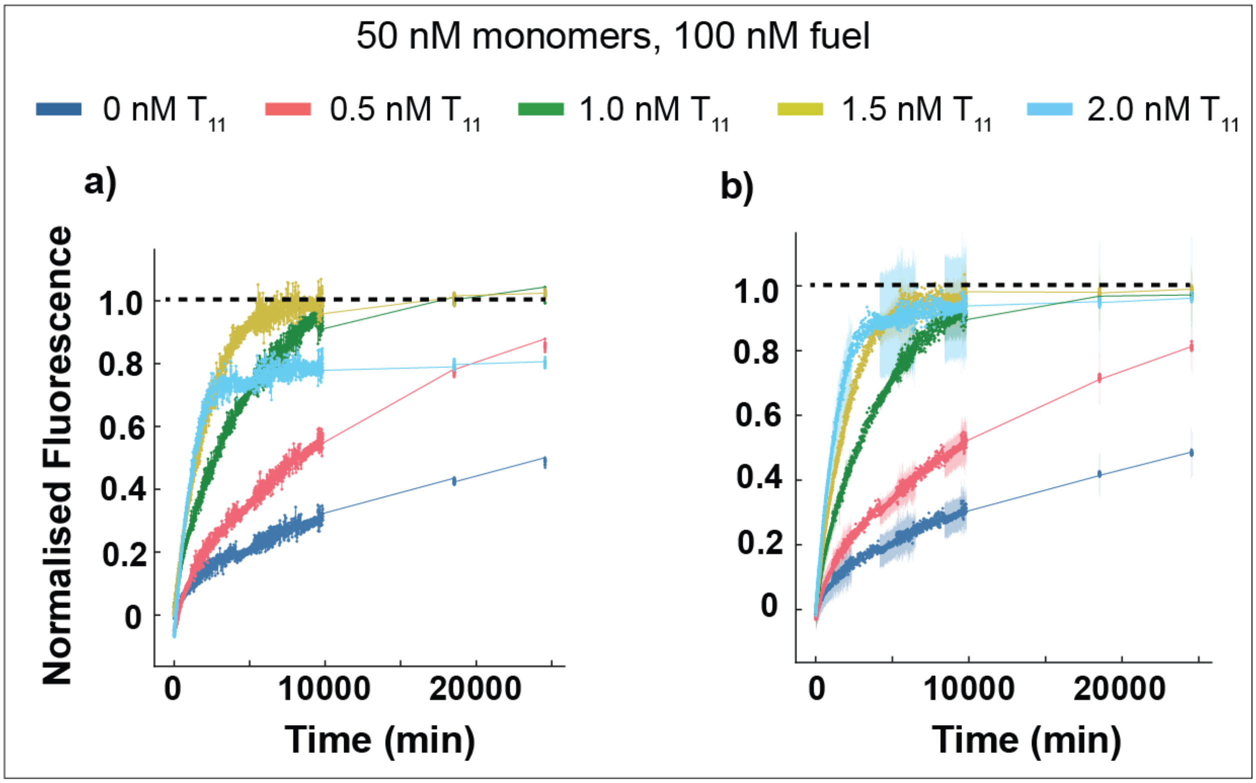

Figure S10. Catalytic templating with high monomer to template concentration ratios. Reactions were conducted according to the experimental protocol outlined in Table S21, with template $T_{11}$ added to $S_l$-L, $R_1$ and $D^7$. The fluorescence in the ATTO 488 channel, normalized with respect 10 nM of $S_1$-$R_1$-$D^7$ according to the procedure in section SI.3.2, is plotted against time. In (a), we show a single replica with anomalous behaviour in the 2.0 nM trace, and in (b) we combine that data with that in Figure 3f, also including subsequent measurements taken at long time.

##### SI.5.6 Reactions for orthogonal dimerization

###### SI.5.6.1 Full reaction for orthogonal set 2

Data from a single replica, showing that the dimerisation of the second set of monomers $S_2$ and $R_2$ can be effectively templated by $T_{22}$ is plotted int S11.

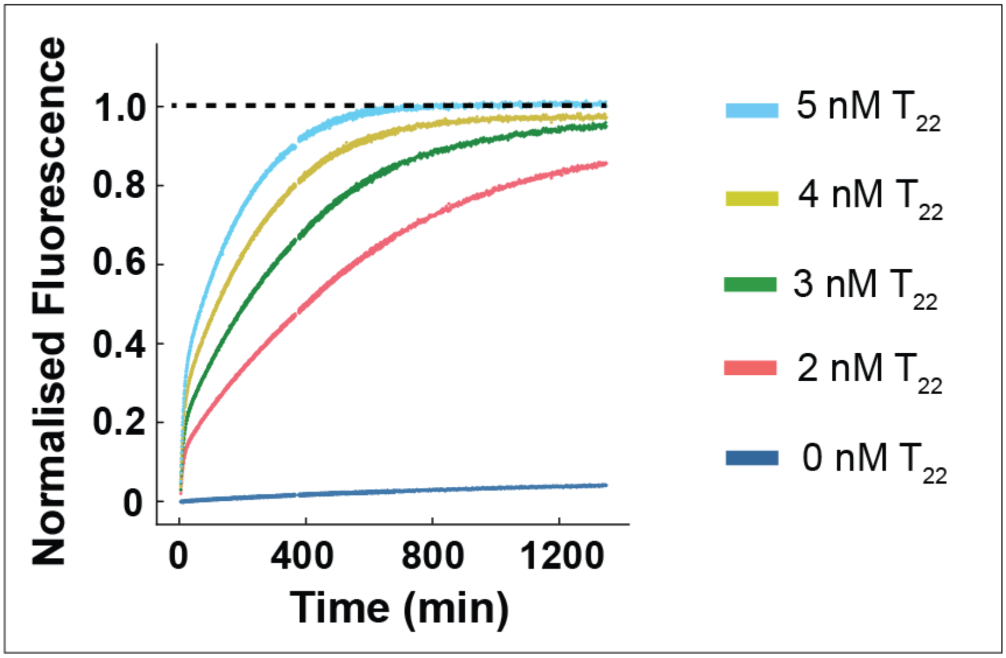

Figure S11. Templating reactions of the second orthogonal set showing catalytic turnover. Reactions were conducted according to the experimental protocol outlined in Table S22, with template $T_{22}$ added to $S_2$-L, $R_2$ and $D^7$. The fluorescence in the Alexa Fluor 546 channel, normalized with respect 10 nM of $S_2$-$R_2$-$D^7$ according to the procedure in section SI.3.2, is plotted against time.

#### SI.5.6.2 Dimerization reactions of each pair of monomers triggered with four different templates

Data complementary to that in Figure 4a in the manuscript showing the behaviour of the other three pairs of monomers, when challenged with 4 different templates, is plotted in Figure S12. All datasets are obtained from single replicas.

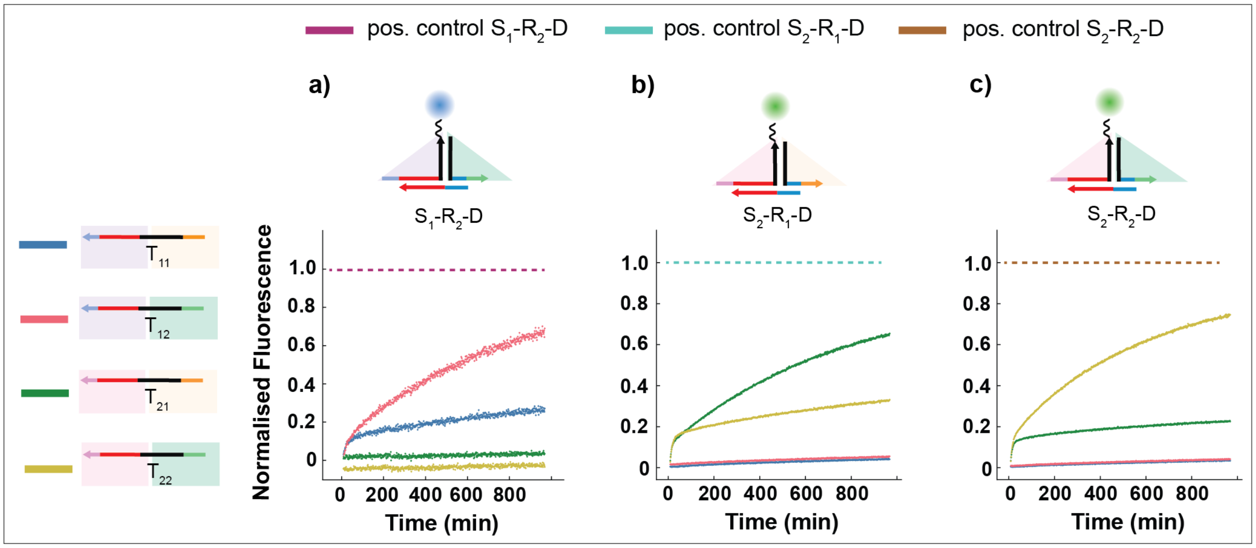

Figure S12. Specificity test for the orthogonal templates with each pair of monomers. Reactions were conducted according to the experimental protocol outlined in Tables S23, with templates $T_{ij}$ separately added to solutions of: $S_1$-L, $R_2$ and $D^7$ in (a); $S_2$-L, $R_1$ and $D^7$ in (b); and $S_2$-L, $R_2$ and $D^7$ in (c). The fluorescence in the ATTO 488 channel (in (a)) or Alexa Fluor 546 channel (in (b) and (c)), normalized with respect 10 nM of the relevant product $S_k$-$R_l$-$D^7$ according to the procedure in section SI.3.2, is plotted against time.

##### SI.5.6.3 Specificity of the templates in a pool of monomers

In Figure S13, we plot for completeness the full gel image corresponding to Figure 4e in the manuscript, following the reaction protocol as described in Table S25.

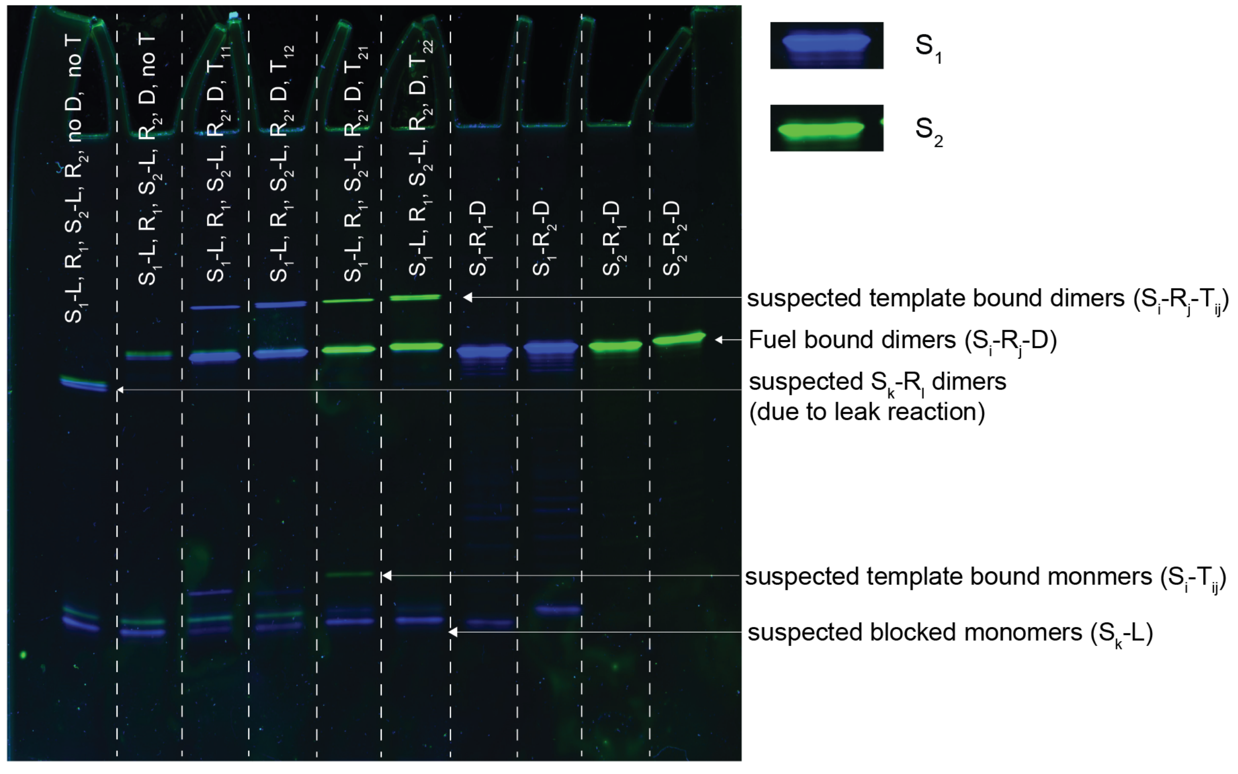

Figure S13. Gel electrophoresis demonstrating specificity of templates and orthogonal dimerization. Reactions were conducted according to the experimental protocol outlined in Table S25, with one of four different templates added to a mixture of all four monomers and fuel D.

##### SI.5.6.4 Gel showing non-reactivity of second monomer and fuel prior reaction

Reaction mixtures were prepared as reported in Figure S14. The reaction mixtures were incubated at 4 °C for 48 h. Then the reaction mixtures and controls were loaded in a PAGE following the protocol in the gel electrophoresis section of the methods in the main manuscript. In this case, the gel was run for 75 min and then imaged on gel scanner. Following this procedure, the gel was stained with SYBR Gold dye and imaged again.

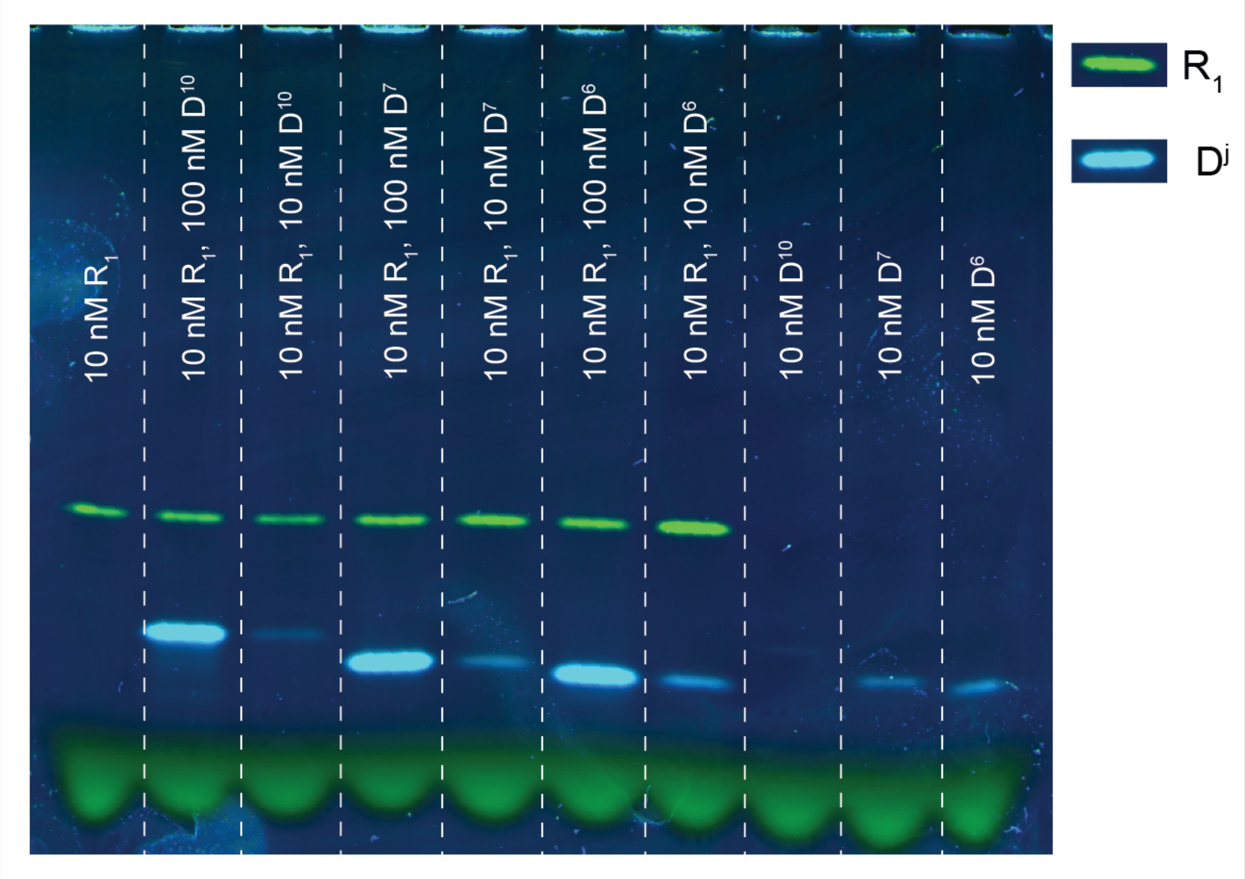

Figure S14. Gel electrophoresis demonstrating no reaction between second monomer and fuel before the triggering the reaction.
